# Global analysis of thermal and chemical denaturation using CheMelt: Thermodynamic dissection of highly thermostable *de novo* designed proteins

**DOI:** 10.64898/2026.04.07.716910

**Authors:** Vili Lampinen, Osvaldo Burastero, Iara Plácido Guazzelli, Florian Vögele, Francisca Pinheiro, Jan S. Nowak, Maria M. Garcia Alai, Magnus Kjaergaard

## Abstract

*De novo* protein design often produces thermostable proteins that denature above 100 °C, which complicates the analysis of their stability. Thermostable proteins can be unfolded by combined chemical and thermal denaturation followed by global analysis of multiple melting curves. Here, we have developed CheMelt, a new online tool for global analysis of unfolding data via an intuitive graphical user interface. We use nanoscale differential scanning fluorimetry followed by CheMelt data analysis to dissect the combined thermal and chemical denaturation of thirty-five *de novo* designed protein binders. Thirteen present sufficient fluorescence changes to extract thermodynamic parameters of unfolding. These *de novo* designed proteins have systematically lower Δ*C*p and *m*-values than comparable natural proteins. We show that a high thermostability of a designed protein does not necessarily imply a high equilibrium stability, and demonstrate the potential of CheMelt in dissecting thermodynamic properties for protein design and engineering. CheMelt can be accessed at https://spc.embl-hamburg.de/app/chemelt

## INTRODUCTION

*De novo* designed proteins often exhibit high thermostability with melting temperatures routinely exceeding the boiling point of water (Cao et al. 2022; Vázquez Torres et al. 2024, 2025; Zheng et al. 2025). Thermostability is attractive for many biotechnological applications and may enhance activity by preserving the folded structure (Pezeshgi Modarres et al. 2016). High thermal stability may even be used for rapid purification of designed proteins by heat lysis (Pinheiro et al. 2025; Grene et al. 2025). While thermostability seems to be a frequent feature of *de novo* designed proteins, its structural and thermodynamic origin is unclear.

Protein thermostability has been extensively studied in the context of evolutionary adaptations for extremely hot environments (Jaenicke and Böhm 1998). Homologous proteins from meso- and thermophiles have been compared to reveal how evolution tunes thermodynamic parameters to achieve thermostability (Sandeep Kumar et al. 2001). Based on these studies, three models of unfolding have been identified, which can be illustrated in a plot of the free energy of unfolding, Δ*G_u_*, as a function of temperature (Figure 1) (Nojima et al. 1977). First, thermophilic proteins could simply be very stable and have a higher Δ*G*_u_ at all temperatures. Second, the stability maximum could be shifted to higher temperatures. This would imply that they are close to cold denaturation at moderate temperatures and are initially stabilized upon heating. Third, the Δ*G*_u_ of thermostable proteins may be less affected by temperature, which is the case when the change in heat capacity upon unfolding (Δ*C*_p_) is small. These mechanisms can work together, and surveys of thermophilic proteins indicate that all three mechanisms are used by naturally thermostable proteins (Razvi and Scholtz 2006).

**Figure 1.**
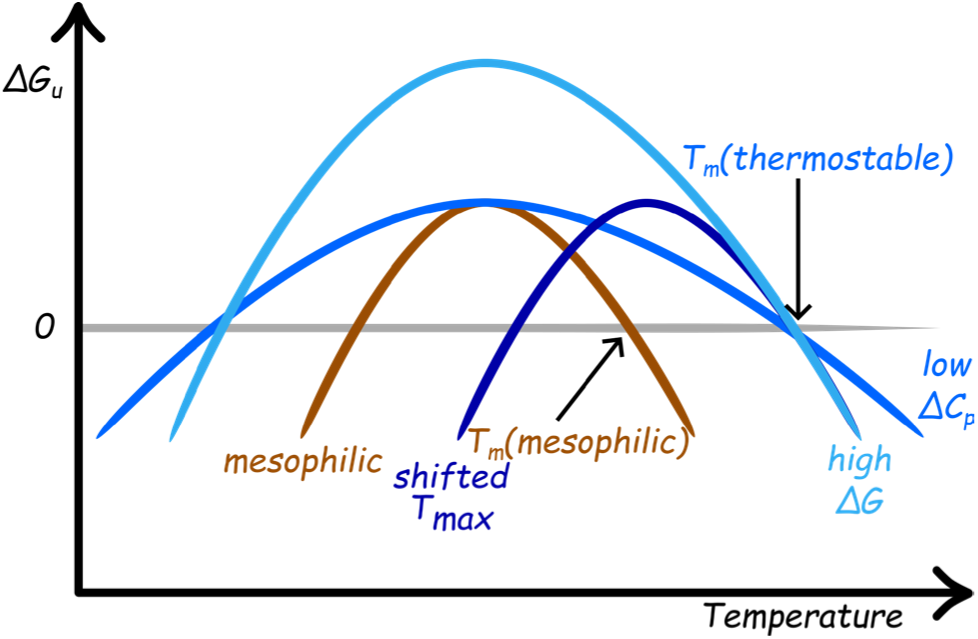
Hypothetical temperature profiles of free energies of unfolding of mesophilic and thermophilic proteins. The melting temperatures for denaturation are obtained from the points on the graph where Δ*Gu*=0. Compared to a similar mesophilic protein (brown), the melting temperature (*T*m) of a thermostable protein can increase in several ways: By an increase in the maximum Δ*Gu* value (light blue), by shifting of an identically shaped curve towards higher temperatures (darkest blue), or by flattening of the curve via a low Δ*C*_p_ (mid-blue).

For protein engineering, thermodynamic parameters of folding need to be determined accurately and with high throughput. Differential scanning fluorimetry (DSF) allows parallel measurement of many samples using minimal protein, so-called nanoDSF, making it highly suited for this purpose (Magnusson et al. 2019; Pantoliano et al. 2001). In a DSF experiment, proteins are thermally denatured while the fluorescence emission of aromatic residues or an extrinsic dye is monitored (Magnusson et al. 2019; Pantoliano et al. 2001). DSF can be extended to thermostable proteins by inclusion of chaotropic denaturants, which lower Δ*G*_u_ linearly with denaturant concentration (Privalov 1979). By recording thermal denaturation traces at different denaturant concentrations, Δ*G*_u_ can be extracted as a function of denaturant concentration and extrapolated to the absence of denaturant. In practice, individual melting curves are often underdetermined, and parameters are more robustly extracted by a global fit of curves recorded at a wide range of denaturant concentrations (Hamborg et al. 2020). Multivariate analysis of 2D unfolding data remains challenging for many laboratories and would benefit from more accessible, standardized analytical tools.

Here, we describe CheMelt, an online user-friendly tool for the global analysis of combined thermal and denaturant-induced unfolding. CheMelt is available at https://spc.embl-hamburg.de/app/chemelt, and is based on the Python package *pychemelt* (https://github.com/osvalB/pychemelt). Here, we first validated CheMelt with a previously published experimental dataset (Case I), then a dataset that simulates a highly thermostable protein (Case II), and finally, we use CheMelt to analyse nanoDSF data from thirteen highly thermostable *de novo* designed minibinders (Case III). We show that these designed proteins mainly achieve high thermostability via lower Δ*C*_p_ values than expected for proteins of their size.

## RESULTS

### SOFTWARE DESCRIPTION

We developed a new web tool called CheMelt for global analysis of denaturant and thermal denaturation. CheMelt is based on previous modules for the analysis of thermal unfolding data only: MoltenProt intended for fitting individual melting curves, and FoldAffinity intended for fitting ligand titration series (Burastero et al. 2021; Niebling et al. 2021). CheMelt is hosted together with a series of other tools for understanding of biophysical data at the eSPC platform (https://spc.embl-hamburg.de/).

The workflow of CheMelt can be conceptually divided into five steps (Figure 2). The first step allows import of files containing temperature, fluorescence intensities, and denaturant concentrations. The server natively supports output files from popular DSF instruments, e.g. Nanotemper Prometheus. In the second step, a fitting model is selected including equations for how the baselines depend on temperature. The intercept and slope terms of the baselines can either be fitted globally (recommended) or individually for each curve. The third step is the actual fitting. For computational efficiency, it is possible to downscale the data density at this stage. The fourth step calls for an evaluation of the fit and parameter quality. Fits that give large and non-random residuals, or high error estimates should be critically re-evaluated. In the fifth step, it is possible to create publication quality figures as showcased in the figures of this manuscript. Practical instructions on how to use the server are detailed in the Supplementary Methods section (SI).

**Figure 2.**
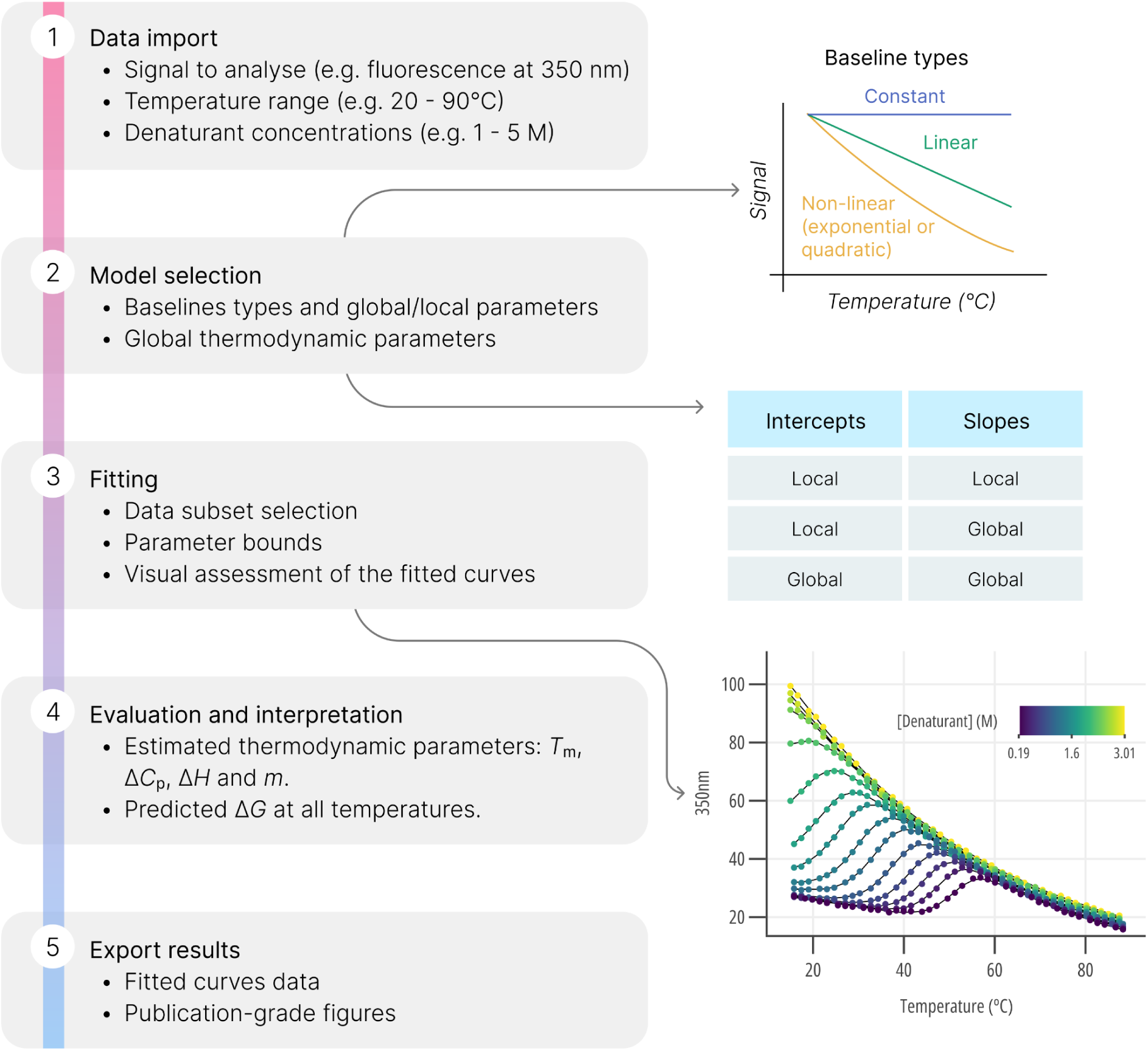
Workflow of CheMelt. 1) Data import, 2) Model selection, including the type of baselines, 3) Global fitting to estimate Δ*C*p, *T*m, Δ*H*m and *m*, 4) Interpretation, and 5) Export of the results.

### CASE I - ALGORITHM VALIDATION USING ACBP EXPERIMENTAL DATA

#### CASE I - ALGORITHM VALIDATION USING REPORTED EXPERIMENTAL DATA

To test CheMelt, we reanalysed two datasets, one from the bovine Acetyl-CoA binding protein (ACBP), a small helical bundle that has been used as a model system for protein folding studies (Teilum, Maki, Kragelund, Poulsen, & Roder, 2002; Thomsen, Kragelund, Teilum, Knudsen, & Poulsen, 2002); and one from the protein DNAJB6b, a chaperone with two folded globular domains connected by a long unstructured region (Fricke et al. 2025).

Previously, the temperature dependence for the native and unfolded states baselines were modelled with a linear and quadratic equation, respectively. However, we found that using an exponential equation for the unfolded baseline improved generalizability by deviating less from a completely unfolded curve after fitting (Figure S1 and S2). Subsequently, an exponential model for the unfolded baseline was used in this analysis. CheMelt derived global thermodynamic parameters, slopes and baselines fit the data well (Figure S3 and S4).

Regarding ACBP, the fitted parameters differ by less than ten percent from the ones previously reported by Hamborg *et al*. (Hamborg et al., 2020) (Table S1). Additionally, both CheMelt and Hamborg *et al*. report *m*-values and free energy of unfolding values that are in close agreement with previous measurements by Teilum *et al*. (Hamborg et al., 2020; Teilum, Kragelund, & Poulsen, 2002). In the case of DNAJB6b, the parameters are consistent between methods, with a difference of less than 10 %, except for Δ*C*_p_ that was fitted with a value of 0.9 kcal/mol/K and was reported to be 0.72 kcal/mol/K (Table S2). This slight difference can be probably attributed to use of exponential baselines and a scale factor in CheMelt, that results in a better fitting with a lower Akaike information criterion and Bayesian information criterion (Table S3). Overall, these results show that CheMelt can fit experimental protein melting data and reproduce previously reported thermodynamic parameters.

### CASE II - SIMULATED DATA OF HYPER-STABLE PROTEINS

A key application for CheMelt is the thermodynamic analysis of highly thermostable proteins. In such data sets, the desired parameters in the absence of denaturant cannot be measured directly, but they rather need to be extrapolated from melting curves recorded at high denaturant concentrations. Before analysing experimental data, we wanted to test whether CheMelt is still able to accurately extract thermodynamic parameters under these conditions using synthetic data. We simulated and fitted unfolding curves with fixed values (similar to ACBP) of Δ*H*_m_, Δ*C*_p_, *m*, but with variable *T*_m_ values spanning from 100 to 140 °C (Figure S5). Our results indicate that, even if there is no visible unfolding transition for the protein without denaturant, it is still possible to estimate a *T*_m_ value, as well as the other thermodynamic parameters (Figure S5).

### CASE III - HYPER-STABLE *DE NOVO* DESIGNED PROTEINS

To explain the origin of the high thermal stability of *de novo* designed proteins, we selected thirty-five protein binders designed using a combination of RFdiffusion (Watson et al. 2023) and proteinMPNN (Dauparas et al. 2022). Seventeen of these proteins were reported previously as putative binders of the GK-domain of PSD-95 and the ALFA tag (Pinheiro et al. 2025). The remaining eighteen were designed as binders against the disordered C-terminus of GluN1 (Table S4). All proteins are predicted to form a compact structure (Figure 4) by AlphaFold2 with a predicted alignment error (pAE) < 6.2 (Jumper et al. 2021) (Table S5), and remain soluble after purification by heat lysis. Except for two proteins (G3A and G5B), all the proteins in our test set were single domain alpha-helical bundles with 96–221 (median 118) amino acid residues.

As expected, none of the binders showed any significant denaturation without the presence of a denaturant, even at temperatures exceeding 90 °C. To identify suitable concentrations of denaturant for unfolding, we acquired nanoDSF measurements for all thirty-five proteins at 0, 2, 4, and 6 M guanidine hydrochloride (GdmCl) (Figure 3). For many of the proteins, a shift in 350 nm/330 nm ratio signal was observed only at one denaturant concentration. Lower and higher concentrations followed the pre- or post-transition baselines suggesting that the protein remains either fully folded or unfolded during the thermal gradient. Surprisingly, twenty out of thirty-five of the tested proteins did not show any transition in either fluorescence intensity at 330 nm, 350 nm, or their ratio (Figure 3C). While all tested proteins have at least one fluorescent residue, the proteins that did show an unfolding transition had more total and buried aromatic residues than those that did not (Figure S6). To test whether these proteins fail to unfold, we measured the unfolding of five proteins with flat nanoDSF traces using far UV circular dichroism spectroscopy (CD). CD showed that two of the proteins were mostly denatured at 25 °C in 6 M GdmCl (Figure S7A), two at 95 °C in 6 M GdmCl, while G15A still retains some ɑ-helix character after spending more than fifteen minutes at 95 °C in 6 M GdmCl (Figure S7B). Therefore, these five proteins all unfold at high denaturant and temperature, but their unfolding does not result in an observable change in fluorescence emission. As the focus of this work is nanoDSF analysis, we did not characterise these proteins further.

**Figure 3.**
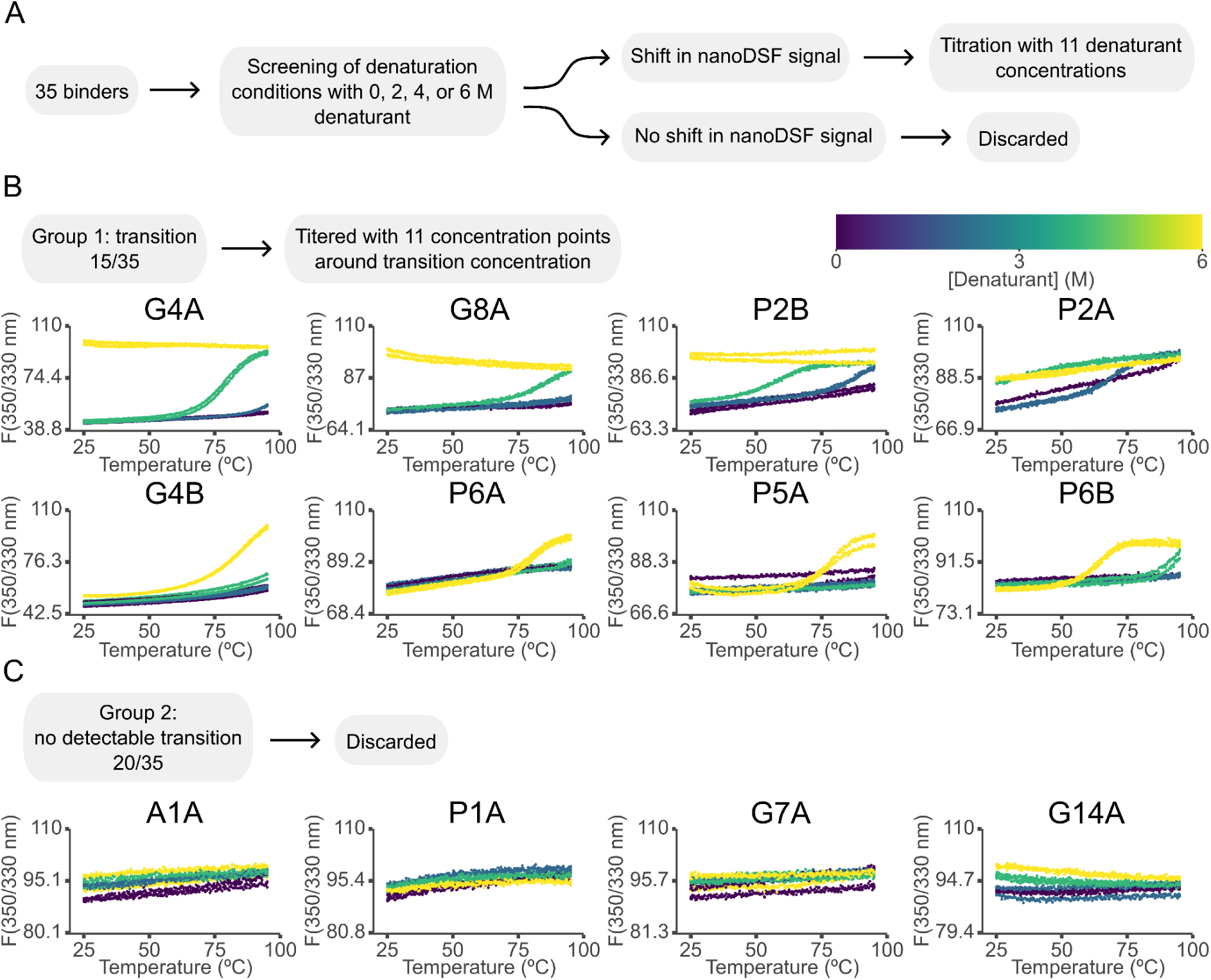
Screening of denaturation conditions for *de novo* designed protein binders. A) Flow chart of the screening process: *De novo* designed proteins were screened by nanoDSF at 0, 2, 4, and 6 M GdmCl to identify proteins with an observable unfolding transition. B) Representative melting curves of proteins that show an unfolding transition at least at one denaturant concentration. C) Representative DSF traces of proteins that do not show an unfolding transition. All graphs show two replicate measurements at each denaturant concentration and were prepared using the built-in tools in the CheMelt web server.

For the fifteen proteins that showed an observable melting transition, we performed nanoDSF at eleven GdmCl concentrations (Figure 4). A4A was included in the titration as it seemed to show a small transition in the screening, but it was subsequently eliminated from further analysis. The concentration range was determined individually for each protein based on the initial screen. Thirteen out of fifteen proteins showed sigmoidal unfolding curves, with a clear post-transition baseline, suggesting that full unfolding was achieved. Many curves show an upward inflection at low temperatures with higher GdmCl concentrations (A4B, G4A, G5A, G8A, P3A, P3B, P5A), indicative of cold denaturation at demanding denaturant conditions. One protein (G4B) failed to fully unfold even at 95 °C in the presence of 6 M GdmCl. All proteins except for G5A, the only beta-sheet containing protein, showed a full refolding upon cooling (Figure S8, S9) justifying the use of a reversible folding model.

**Figure 4.**
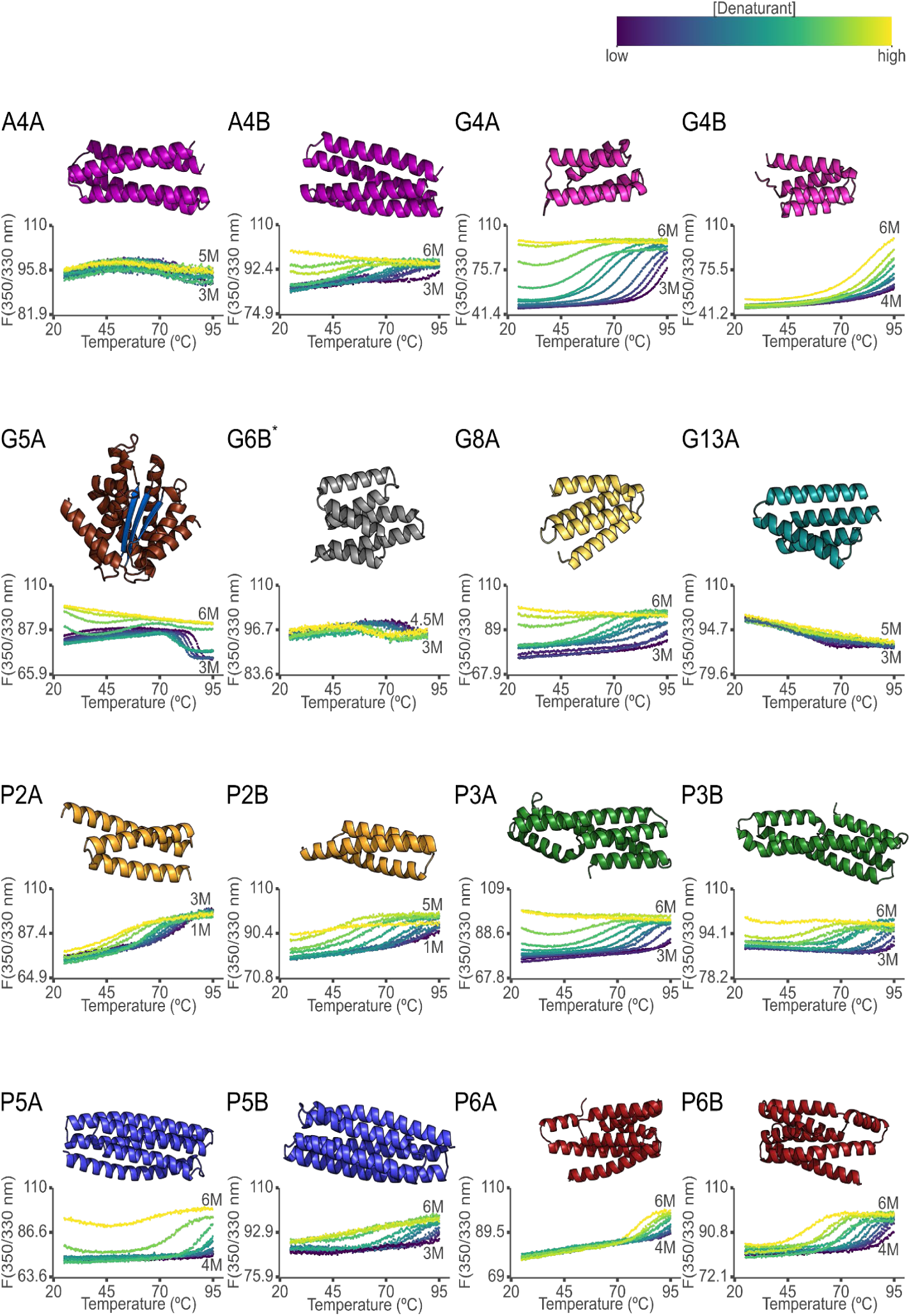
Combined thermal and denaturant titration of *de novo* designed proteins. Proteins with a detectable unfolding transition were titered in 0.2 or 0.3 M steps around the denaturant concentrations showing a transition in the screening. The highest (yellow) and lowest (purple) denaturant concentration is shown within each graph. Protein models represent an AlphaFold2 prediction above the corresponding melting curves and were prepared with PyMOL 3.1.1 (Schrödinger, LLC 2024). Proteins with the same fold, but different sequences are shown with the same colors. A4A was included based on a minor shift during the screening, but was excluded from further analysis after the titration. For clarity, only a few concentrations are shown for protein G6B. All graphs were prepared using the built-in tools in CheMelt web server.

The ratio of fluorescence intensity (350 nm / 330 nm) is useful for visual inspection of melting curves; but it is not guaranteed to be proportional to the population of molecular species (Žoldák et al. 2017). Therefore, we used the raw fluorescence intensities for globally fitting all eleven melting curves for each protein (Figure 5). We used the 330 nm signal as it was most reproducible between repeats. The different appearances of the fluorescence traces necessitate different baseline models, so we tried progressively more complex baseline models for each protein until we found one that resulted in a good fit. The fit was regarded as successfully converging when the residuals were small, the relative error of the fitted parameters was less than 5%, and the thermodynamic parameters were plausible.

**Figure 5.**
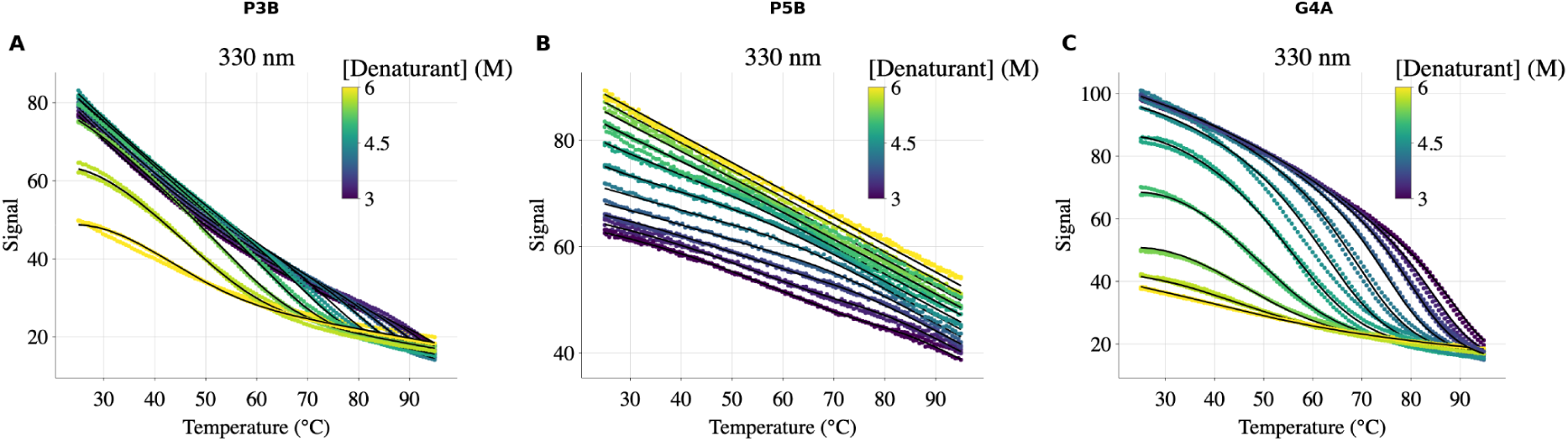
Fitting of denaturation data for *de novo* binder proteins. The model was fitted to the 330 nm fluorescence intensity signal for each titered protein. The figure shows the fitted curves of P5B, G4A and P3B after normalisation by the fitted scaled factor to improve clarity. Scatter plots show the experimental data, with color coding (viridis scale) indicating the GdmCl concentration. Solid black lines represent the global fits to the unfolding model at each denaturant concentration.

CheMelt produced robust fits to thirteen proteins (Fig. S10) with thermodynamic parameters reported in Table I. As expected, all proteins were highly thermostable with melting temperatures near or above the boiling point of water (93 - 173 °C). The highest calculated melting temperature was 173 °C for P5A and the lowest 93 °C for P2A, with an average of 125 °C. The predicted stability curves are shown in Figure S11.

**Table I.**
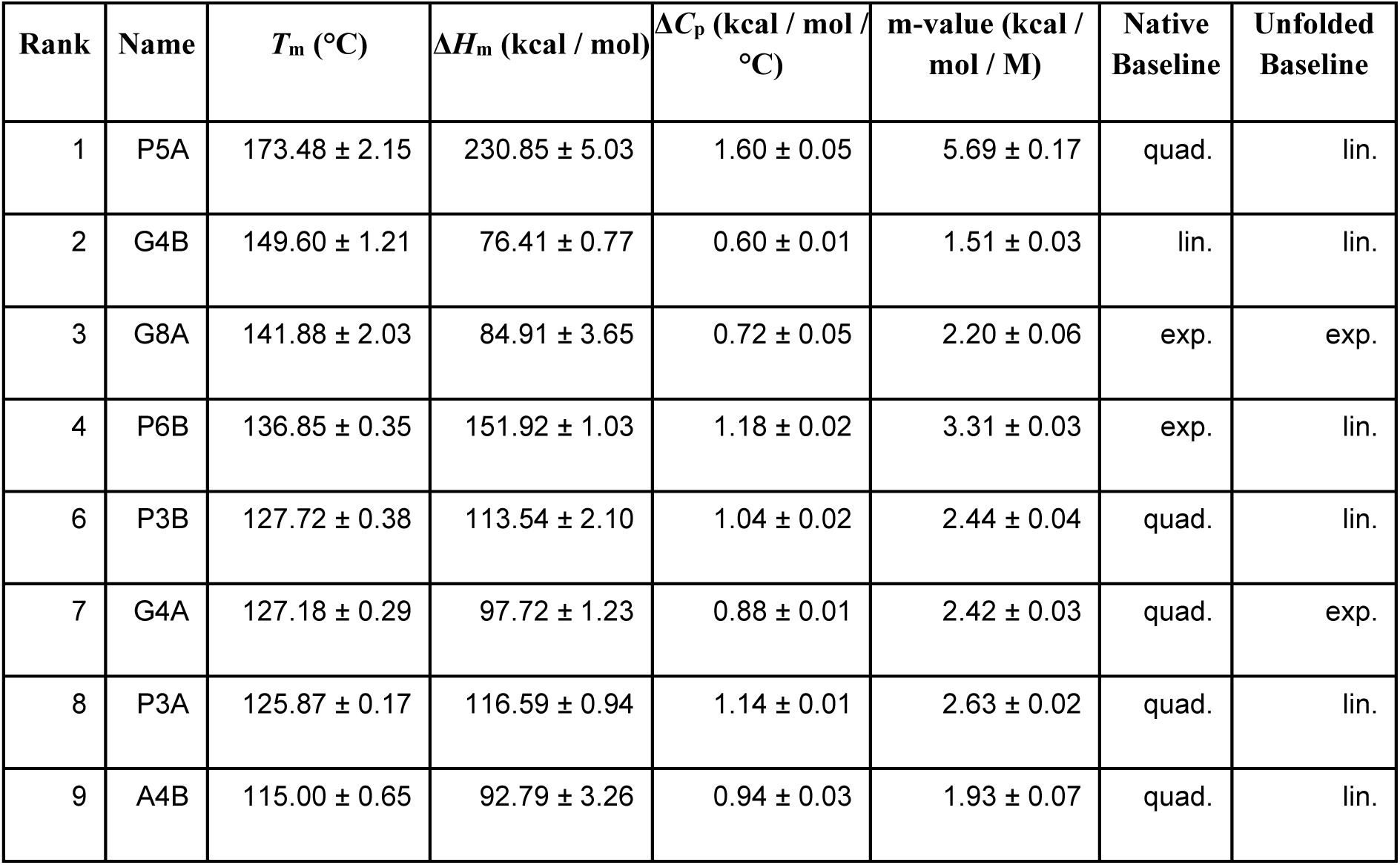
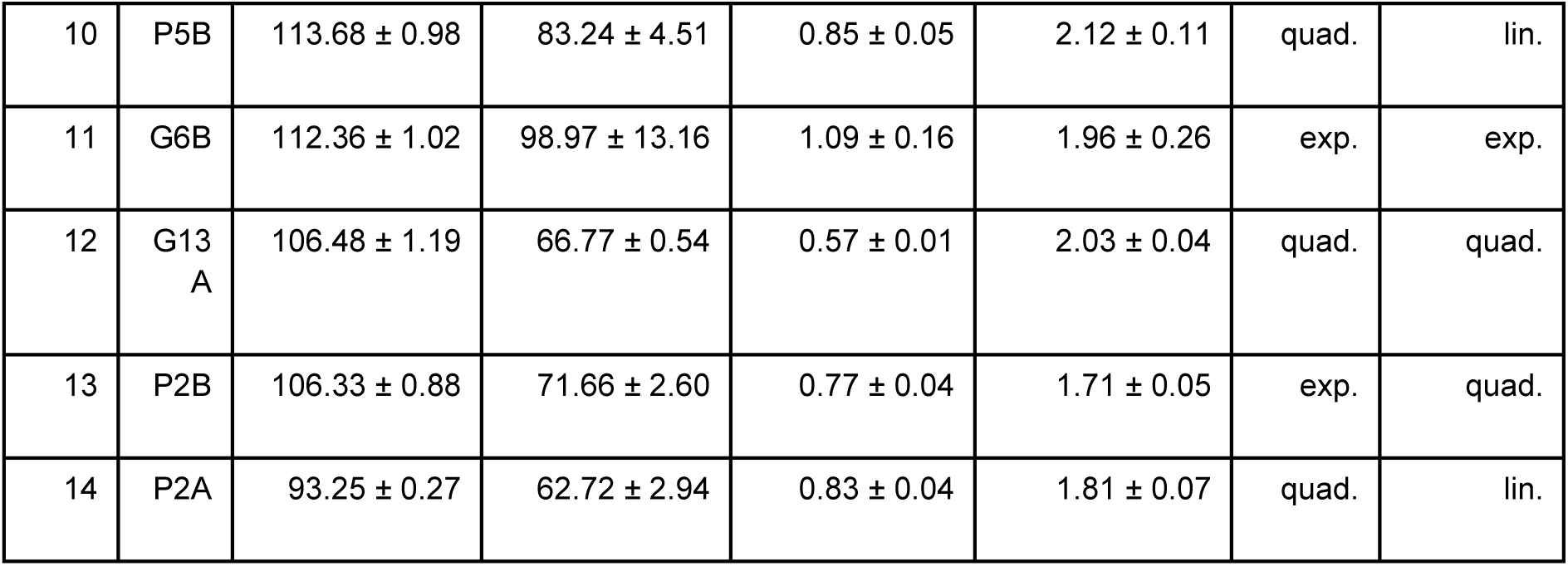
Calculated thermodynamic parameters for the titered *de novo* binder proteins. The proteins are ranked by melting temperature, from highest to lowest *T*m. The reported values represent the mean and confidence intervals calculated using the leave-one-out method. The baselines refer to the baseline model that gave the best fit as indicated by the extended Bayesian Information Criterion.

| Rank | Name | $T_m$ (°C) | $\Delta H_m$ (kcal / mol) | $\Delta C_p$ (kcal / mol / °C) | m-value (kcal / mol / M) | Native Baseline | Unfolded Baseline |
| --- | --- | --- | --- | --- | --- | --- | --- |
| 1 | P5A | 173.48 ± 2.15 | 230.85 ± 5.03 | 1.60 ± 0.05 | 5.69 ± 0.17 | quad. | lin. |
| 2 | G4B | 149.60 ± 1.21 | 76.41 ± 0.77 | 0.60 ± 0.01 | 1.51 ± 0.03 | lin. | lin. |
| 3 | G8A | 141.88 ± 2.03 | 84.91 ± 3.65 | 0.72 ± 0.05 | 2.20 ± 0.06 | exp. | exp. |
| 4 | P6B | 136.85 ± 0.35 | 151.92 ± 1.03 | 1.18 ± 0.02 | 3.31 ± 0.03 | exp. | lin. |
| 6 | P3B | 127.72 ± 0.38 | 113.54 ± 2.10 | 1.04 ± 0.02 | 2.44 ± 0.04 | quad. | lin. |
| 7 | G4A | 127.18 ± 0.29 | 97.72 ± 1.23 | 0.88 ± 0.01 | 2.42 ± 0.03 | quad. | exp. |
| 8 | P3A | 125.87 ± 0.17 | 116.59 ± 0.94 | 1.14 ± 0.01 | 2.63 ± 0.02 | quad. | lin. |
| 9 | A4B | 115.00 ± 0.65 | 92.79 ± 3.26 | 0.94 ± 0.03 | 1.93 ± 0.07 | quad. | lin. |
| 10 | P5B | $113.68 \pm 0.98$ | $83.24 \pm 4.51$ | $0.85 \pm 0.05$ | $2.12 \pm 0.11$ | quad. | lin. |
| 11 | G6B | $112.36 \pm 1.02$ | $98.97 \pm 13.16$ | $1.09 \pm 0.16$ | $1.96 \pm 0.26$ | exp. | exp. |
| 12 | G13<br>A | $106.48 \pm 1.19$ | $66.77 \pm 0.54$ | $0.57 \pm 0.01$ | $2.03 \pm 0.04$ | quad. | quad. |
| 13 | P2B | $106.33 \pm 0.88$ | $71.66 \pm 2.60$ | $0.77 \pm 0.04$ | $1.71 \pm 0.05$ | exp. | quad. |
| 14 | P2A | $93.25 \pm 0.27$ | $62.72 \pm 2.94$ | $0.83 \pm 0.04$ | $1.81 \pm 0.07$ | quad. | lin. |

Protein unfolding leads to a change in the solvent accessible surface area (ΔSASA), which is tightly correlated to the heat capacity change upon unfolding (Δ*C*_p_) and the dependence of the free energy on the denaturant concentration (*m*-value) (Myers et al. 1995). As ΔSASA scales with protein chain length, both Δ*C*_p_ and *m*-values scale linearly with the number of residues, and the two parameters are highly correlated. We compared the Δ*C*_p_ and *m*-values from our present data set of *de novo* designed proteins with a dataset collected from natural single domain proteins from Myers *et al*. 1995 (Myers et al. 1995). The Δ*C*_p_ and *m*-values in our dataset exhibit a correlation and slope similar to those reported by Myers *et al*. indicating internal consistency of the data and suggesting that they reflect ΔSASA upon unfolding (Figure 6A). However, the correlation of Δ*C*_p_ and *m*-value with the number of residues showed a substantially lower slope than found by Myers *et al..* This could in part be because the set of designed proteins mainly consists of elongated helical bundles (as reflected in their higher aspect ratio (Figure S12A)) that expose less surface area per residue upon unfolding. Consistent with this, ΔSASA per residue was significantly lower for the designed proteins (Figure S12B– C), suggesting that their non-globular folds contribute to their low ΔC_p_. However, the ΔC_p_ values of *de novo* designed proteins are lower than for natural proteins that exposed the same amount of surface area by about a third (Figure 6B), and a similar trend is seen for m-values (Figure 6C) suggesting that this is not just an effect of the fold.

**Figure 6.**
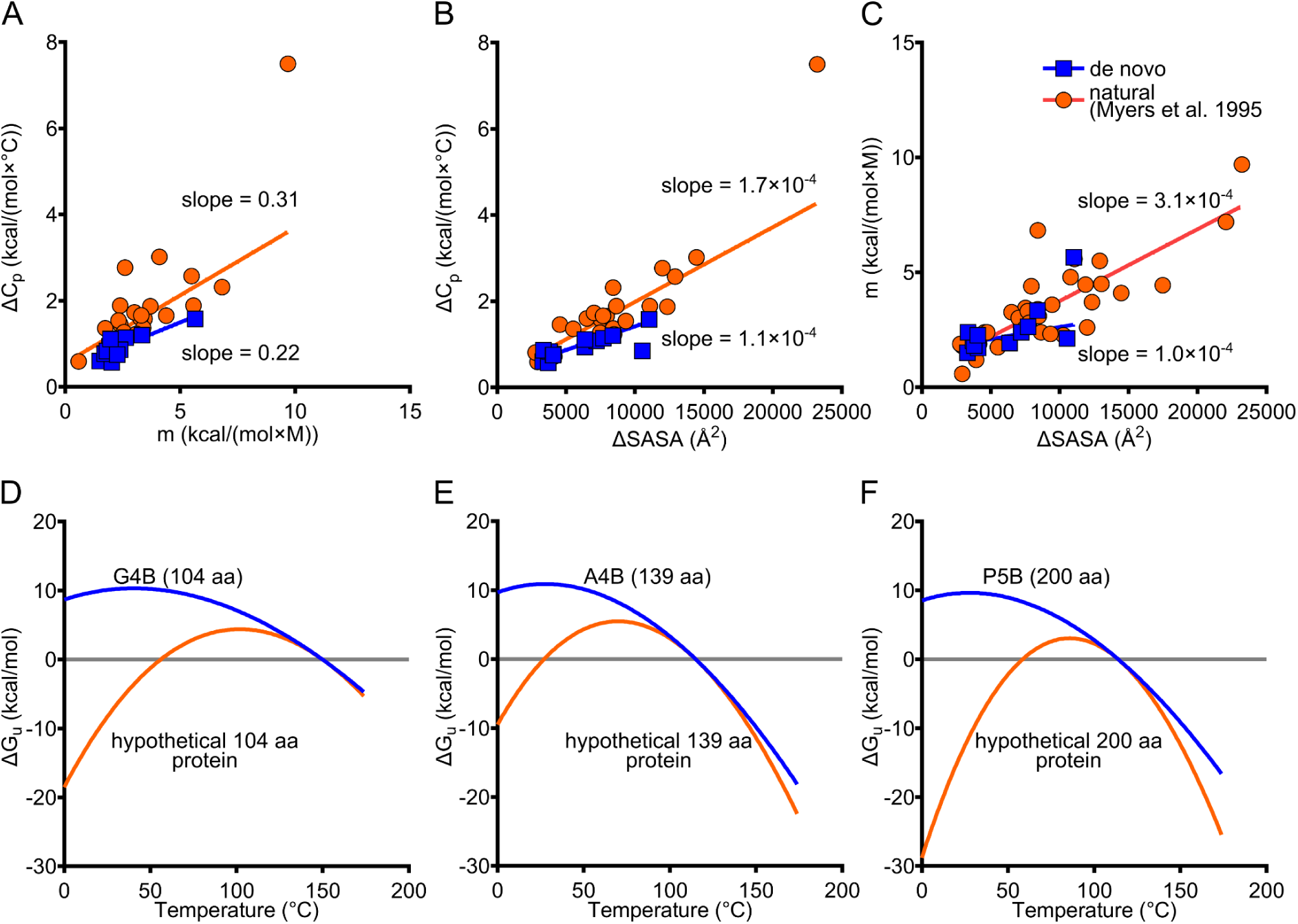
Comparison of thermodynamic parameters of *de novo* binders and native proteins. The list of native proteins and their thermodynamic parameters were extracted from Myers *et al*. 1995 (Myers et al. 1995). B–C) Change in surface accessible surface area upon unfolding (ΔSASA) was estimated using the minimum (compare to Figure S13 for maximum) area per residue in the unfolded state (Creamer et al. 1997) and the Shrake–Rupley algorithm for the folded model (Shrake and Rupley 1973). The lines show linear regression of the data. D–F) The stability plots of three example proteins from the study are compared with proteins with the same length, melting temperature and *ΔH*_m_*(Tm)* value but with a predicted *ΔC_p_* value, instead of an experimentally determined one. The plots were prepared using GraphPad Prism 11.0.0. The predicted Δ*C*_p_ value was calculated as 0.0142 kcal/mol\**n* + 0.0025 kcal/mol, where *n* is the number of residues (Myers et al. 1995).

The lower ΔSASA of the designed proteins might in part be explained by a higher degree of structure in the unfolded state. Our designed proteins have a high intrinsic helix propensity as estimated by ABSEIL (Hortman et al. 2026) (Figure S13), which might lead to a higher residual structure in the unfolded state. Estimation of ΔSASA relies on subtraction of the accessible surface area of the unfolded state, which cannot be measured directly, but has to be approximated. In principle, the surface area of the unfolded state can be estimated more precisely from the radius of gyration (Geierhaas et al. 2007), but this is not experimentally accessible due to the harsh conditions needed to unfold the designed proteins. Creamer et al. (Creamer et al. 1997) modelled the effect of residual structure in the unfolded state by calculating the exposed surface area of tri-peptides from folded and extended chains and thus providing an upper and a lower bound on the accessible surface area in the unfolded state. Figure 6B uses the value of the lower bound for accessible surface area in the unfolded state and thus the lower bound estimate of ΔSASA. Using the upper bound for accessible surface area (Figure S14) does not change the outcome. In combination, this suggests that neither difference in the fold nor in the unfolded state is sufficient to describe the difference between designed and natural proteins.

In the stability plots in Figure 1, Δ*C*_p_ determines the curvature as the second derivative of the free energy of folding with respect to the temperature. To assess how our *de novo* proteins behave relative to natural proteins, we compare stability plots derived from experimental parameters (Δ*C*_p_, Δ*H (T*_m_), and *T*_m_) with hypothetical plot constructed using the Δ*C*_p_ value expected for a protein of the same length (Myers et al. 1995) (Figure 6D-F). The *de novo* designed proteins have a flattened stability plot, with the effect being most pronounced for larger proteins. Incidentally, small proteins always have a relatively low thermal sensitivity due to their small hydrophobic cores (Alexander et al. 1992). In conclusion, the proteins in our data set primarily achieve their high thermostability through a reduced temperature sensitivity rather than a high equilibrium stability, as has been previously proposed for thermophilic proteins (Zhou 2002).

## DISCUSSION

We have developed CheMelt as a graphical user interface for global analysis of combined denaturant and thermal denaturation of proteins. By providing a user-friendly, web-based server, we aim to lower the barrier to entry and promote broader use of this methodology. The ability to determine thermodynamic parameters was similar to that reported in (Hamborg et al. 2020).

The rise of *de novo* protein design will increase the need for analysis of thermostable proteins. Nanoscale DSF is a convenient candidate for such studies, although many recent design papers have used CD spectroscopy to assess stability e.g. (Cao et al. 2022; Vázquez Torres et al. 2024, 2025; Koga et al. 2020; Zheng et al. 2025). Compared to CD, nanoDSF uses less material and is easier to run for many samples in parallel. A disadvantage is that not all proteins will have a detectable unfolding titration in the fluorescence emission. Here, we only observed an unfolding transition for 15 out of 35 (43%) of the proteins tested. Using CD spectroscopy, we showed that a selection of proteins without a fluorescence transition unfold with similar melting temperatures as the ones with a detectable transition. This suggests a minimal environmental change of the aromatic residues upon unfolding, pointing to a difference in the hydrophobic cores of the designed proteins. Our test set of designed proteins is dominated by small helical bundles, which do not leave much room for bulky tryptophan residues in the interior, especially in the idealized tight packing favored by machine learning algorithms. The lack of fluorescence changes may generalise to other helical mini-binders, but not necessarily to designed proteins with other folds.

Fitting thermodynamic models to fluorescence-based unfolding data will often be an ill-posed problem, so a set of unfolding curves can be explained by different combinations of parameters. Therefore, it is recommended to use a broad range of denaturant concentrations to adequately constrain the parameter space. Even then, the fitting procedure may yield physically unrealistic parameter values if the optimization routine is trapped in a local minimum. Compared to other experimental techniques, nanoDSF data is challenged by pre- and post-transition baselines that are often non-linear. The fluorescence emission of indole compounds, which are models for tryptophan, have a non-linear temperature dependence that is modulated by the solvent (Kirby and Steiner 1970; Klein and Tatischeff 1977). It is thus likely that different proteins will require different baseline models observed in our data set (Table 1).

Compared to melting curves for protein folding model systems (e.g. ACBP), the designed proteins tested here are more challenging to fit due to the combination of a small change in fluorescence parameters as discussed above, and a broad unfolding transition. The latter is a consequence of the low Δ*C*_p_ and shallow dependence of equilibrium constant on temperature discussed below. From a fitting perspective, this makes it more difficult to deconvolute the sloping baselines and the unfolding transition. Even under these challenging conditions, CheMelt converges to unique parameter sets for the thirteen proteins tested here. It should be noted that the fitting routine is not guaranteed to work for all proteins, and especially prone to fail if the data set cannot uniquely constrain the baselines in at least one melting curve. For highly thermostable proteins, it should be ensured that the highest denaturant concentration reveals sufficient post-transition baseline before fitting.

We found that the Δ*C*_p_ value of unfolding of the *de novo* designed proteins is much lower than for natural proteins of a similar length. As far as possible, we have excluded error sources that could lead to this conclusion by testing the robustness of the global fits to baseline models and initial parameters. We have not been able to exclude a potential contribution from a hypothetical temperature dependence of the m-values, although they are almost universally assumed to be temperature independent. However, such independence is not strictly guaranteed when the unfolded state has temperature-dependent residual structure. Such an effect would likely be less than the factor of 2-3 we see for the *de novo* designed proteins. The low Δ*C*_p_ values do not necessarily generalize to all *de novo* designed proteins as a few caveats should be noted: The dataset used here are all mini-binders with exposed hydrophobic binding patches, and all have been designed with a similar workflow based on RFdiffusion (Watson et al. 2023) and proteinMPNN (Dauparas et al. 2022) followed by ranking by AlphaFold2 (Jumper et al. 2021). Similarly, the data set is almost exclusively composed of small ɑ-helical bundles, and the only β-sheet containing protein was a noticeable outlier. Nevertheless, our results show that a low Δ*C*_p_ is one mechanism by which protein design algorithms can achieve thermostability, similar to what has been proposed for proteins from thermophilic organisms (Razvi and Scholtz 2006).

Large positive Δ*C*_p_ values for protein folding are widely accepted to reflect the burial of hydrophobic groups, in analogy with the effect observed upon transfer of hydrocarbons to water (Spolar et al. 1992; Baldwin 1986). Alternatively, *de novo* designed proteins may be systematically different from natural proteins and this impacts their folding thermodynamics. A recent NMR survey suggested that many *de novo* designed proteins exhibited regions with non-uniform dynamics, despite being designed to form a static structure (Müntener et al. 2026). In-depth folding studies of a designed protein thus suggested that its folding was less cooperative than natural proteins of similar size (Watters et al. 2007), which simulations attributed to a more frustrated energy landscape (Neelamraju et al. 2018), and mutations of the hydrophobic core of a hyperstable *de novo* designed protein resulted in a more dynamic core and a lower Δ*C*_p_ without changing the overall fold (Koga et al. 2020). While our present data set is neither large enough nor general enough to deduce the origin of the low Δ*C*_p,_ it is consistent with previous findings that suggest that the folded states of these kinds of *de novo* designed binder proteins may be more dynamic.

High stability is often highlighted as an advantage of *de novo* designed proteins. In certain applications, thermal stability is an essential property, for example in industrial processes that occur at high temperatures. In other cases, high thermal stability may be viewed as a proxy for stability towards other types of perturbations. In the latter case, it is important to distinguish between high thermal stability achieved through a high Δ*G*_u_, and through a low Δ*C*_p_. A high Δ*G*_u_ implies a low fraction of the unfolded state at room temperature. Therefore, a high Δ*G*_u_ will likely also translate into robustness against undesirable reactions occurring via the unfolded state such as aggregation or proteolysis. High thermal stability achieved predominantly through a low Δ*C*_p_ does not imply a low fraction of unfolded state at room temperature. By extension, thermostability achieved through a low Δ*C*_p_ will likely not provide robustness against other challenges of the integrity of the protein. When characterising the stability of proteins in design and engineering, it is thus advisable to do a full characterisation of the thermodynamics parameters rather than relying on melting temperature as the sole metric for stability. CheMelt provides a robust data analysis platform for such characterisation.

## MATERIALS AND METHODS

### SIMULATED DATA

The Case II free energy of unfolding was calculated using these parameters: Δ*H*_m_ (100 kcal·mol^−1^), Δ*C*_p_ (1 kcal·mol^−1^·K^−1^), and *m* (3 kcal·mol^−1^·M^−1^). The *T*_m_ values ranged from 70 °C to 120 °C at 10 °C intervals, totaling six datasets. The folded state baseline had a linear dependence on temperature and denaturant concentration, while the unfolded state baseline had an exponential dependence on temperature and a linear dependence on denaturant concentration:

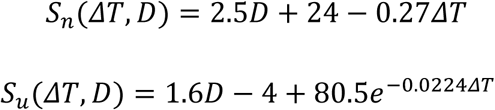

where *S_n_* is the native state baseline and *S_u_* is the denatured state baseline. *ΔT* is the difference between the temperature and a reference temperature (298.15 K) and *D* is the denaturant concentration in molar units.

For each simulated dataset, the temperature ramp went from 20 to 90 °C using steps of 0.7 °C. The denaturant concentration went from 1 to 5, using steps of 0.5 M, resulting in eight unfolding curves per dataset. To mimic experimental conditions, Gaussian noise with a standard deviation of 0.25 was added to each data point, and each unfolding curve was multiplied by a random number between 0.9 and 1.1 drawn from an uniform distribution.

### EXPERIMENTAL DATA

#### ACBP fitting

The unfolding curves of ACBP were downloaded from Github (https://github.com/KULL-Centre/ProteinUnfolding2D/tree/main/examples/data) (Hamborg et al. 2020). The sixteen unfolding curves were measured between 15 and 95 °C, at different GdmCl concentrations up to 3 M in approximately 0.2 M steps. The zero molar denaturant concentration was excluded from the analysis. The signals at 330 nm and 350 nm were fitted simultaneously with *pychemelt*. A linear baseline was used for the native state and an exponential baseline for the unfolded state. The data were reduced to a subset of maximally 200 data points per curve prior to fitting. The model was fitted to the data with globally shared thermodynamic parameters, slopes and baselines. A scaling factor was applied to each concentration to correct for experimental variability. Parameter errors were estimated using the profile likelihood method.

#### DNA6Jb fitting

The unfolding curves of DNA6Jb were downloaded from DTU DATA (https://data.dtu.dk/articles/code/Fitting_of_thermal_and_chemical_denaturation/27679716) (Fricke et al. 2025). The eighteen unfolding curves were measured between 20 and 95 °C, at different GdmCl concentrations up to 6.4 M in approximately 0.4 M steps. The zero molar denaturant concentration, and concentrations higher than 4, M were excluded from the analysis. The signals at 330 nm and 350 nm were fitted simultaneously with *pychemelt*. A linear baseline was used for the native state and an exponential baseline for the unfolded state. The model was fitted to the data with globally shared thermodynamic parameters, slopes and baselines. A scaling factor was applied to each concentration to correct for experimental variability. Parameter errors were estimated using the profile likelihood method.

#### Design and production of proteins

The *de novo* binder proteins (targeting ALFA-tag, GluN1 C terminal domain (CTD), PSD-95 GK-domain) were designed using RFDiffusion (Watson et al. 2023) and ProteinMPNN (Dauparas et al. 2022). The first letter of the name of each binder refers to the binding target, whereas the latter letter indicates different sequences with the same fold. A and P minibinders were reported previously. G minibinders (Table S4) were designed against a peptide (VLPRRAIEREEGQLQL) from GluN1 CTD and selected by a low PAE_interaction_ as predicted by AlphaFold2, which also confidently predicts their monomeric structure (Table S5). The binders were ordered as synthetic genes from Twist Bioscience (USA), who also subcloned them into pET-28a(+) vectors using the NdeI/XhoI restriction sites, so all binders have a 6×His-tag followed by a thrombin site (GluN1 binder sequence Table S4). All binders were expressed in *E. coli,* heat-lysed at 95 °C and purified with Ni-NTA chromatography as previously described (Pinheiro et al. 2025). The proteins were flash-frozen and stored at −20 °C until analysis. All binders were more than 95% pure after the chromatography based on visual inspection of SDS-PAGE gels (Figure S15, other binders described earlier (Pinheiro et al. 2025)). Binders G5A, G14A, G10A, and G4A showed as double bands in the gel (with the smaller band assumed to be a product of protolysis of the binder), but they were still included in the screening stage, and G5A and G4A into titration.

#### Nanoscale differential scanning fluorimetry screening

Unfolding was initially screened for thirty-five binders based on their intrinsic fluorescence with nanoDSF (Prometheus Panta, Nanotemper, Germany). Each binder was measured at a final concentration of 10 µM in the presence of 0, 2, 4, or 6 M GdmCl dissolved in phosphate-buffered saline (PBS, pH 7.4). Two replicates of each sample were prepared in 10 µL capillaries (Prometheus standard capillaries, Nanotemper, #PR-C002), and the capillaries were heated in the instrument from 25 to 95 °C, 4 degrees/min with no refolding observation or dynamic light scattering (DLS) measurement.

#### Nanoscale differential scanning fluorimetry titration

For GdmCl titration, two replicate samples of the binders were prepared as described above with eleven GdmCl concentrations around the concentration showing a transition in the screening. We titered all sixteen binder hits in five different concentration groups with increment increases of 0.2 or 0.3 M GdmCl that span a range of 2 or 3 M change in concentration. In the titration unfolding measurement, the temperature was increased one degree/min from 25 to 95 °C and cooled back to 25 °C at the same rate to measure refolding.

#### Fitting of nanoDSF data of *de novo* designed proteins

NanoDSF data was transferred into pyCheMelt by exporting the unfolding curves as a raw data Excel file using the Nanotemper Panta Analysis software. The 330 nm fluorescence intensity was fitted to a two-state unfolding model as described above for ACBP data (global slopes and global intercepts with scale factor). Several possible combinations of baseline types (linear, quadratic, exponential) were tested for the native and unfolded states, and the selected model (Table S6) was the one with the lowest extended Bayesian Information Criterion (EBIC) using a gamma penalty of one. Two proteins were left out of the analysis: G5A and P2A. In the case of G5A, the fitted curves did not match the data well, and the protein did not refold upon cooling, which is a prerequisite for using the model. For P6A, the estimated ΔC_p_ was highly dependent on the baseline choice, while in all the other cases the first and second ranked models reported similar values. The thermodynamic parameters and associated errors were estimated using the leave-one-out method (by fitting the data multiple times leaving one of the unfolding curves at a time).

For comparing *de novo* binders to native proteins, the change in surface accessible surface area upon unfolding (ΔSASA) was estimated using the minimum area per residue in the unfolded state (Creamer et al., 1997) and the Shrake–Rupley algorithm for the folded model (Shrake & Rupley, 1973). This was done for both protein groups for comparability, since the ΔSASA values in Myers et al., 1997 are overestimated with an older model (Creamer et al. 1997).

#### Circular dichroism spectroscopy (CD)

Five *de novo* designed proteins (G12A, A1A, G10B, A1B, and G15A) that did not show a fluorescence shift were studied by circular dichroism (CD). We measured the CD spectra on a Jasco-810 spectropolarimeter (Jasco, Tokyo, Japan) at 25 °C using a thermoelectric temperature controller (Jasco-PTC-348W1). The CD measurements were made in a 1 mm quartz cuvette using wavelengths between 190 and 260 nm with a data pitch of 1 nm, scanning speed of 50 nm/min and a 1 nm bandwidth. For some CD measurements, we did only two measurements and show a representative one, for others, we show averages of three replicates. All CD spectra were background corrected by subtracting the signal of the relevant buffer. We truncated the spectra to start at 212 nm to cut out the part where the high-tension voltage on the photomultiplier tube exceeds 650 V due to GdmCl signal. The data was analysed using the ChiraKit web tool (Burastero et al. 2025).

### THEORY

#### COMBINED CHEMICAL AND THERMAL DENATURATION

The combination of both denaturation methods results in the following expression for the free energy of unfolding (See Supplementary Methods section, SI):

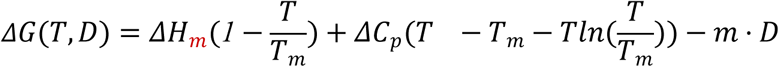

where *T* is the temperature in Kelvin units, *D* is the denaturant concentration in molar units, Δ*H*_m_ is the enthalpy of unfolding, *T*_m_ is the temperature of melting, Δ*C*_p_ is the heat capacity of unfolding, and *m* is an empirical parameter that allows modelling Δ*G* with a linear dependence on denaturant concentration (Greene and Pace 1974). The fraction of protein in the folded (*f*_n_) and unfolded (*f*_u_) states is then calculated as:

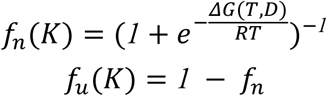

where *R* is the gas constant.

#### FITTING EQUATION

The total signal is assumed to be a linear combination of the signal produced by the different protein states weighted by their mole fractions.

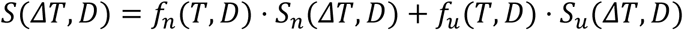

where *S*_n_(*T*, *D*) and *S*_u_(*T*, *D*) are functions describing the signal dependence on temperature and denaturant concentration for the native and unfolded states, respectively. The dependence on temperature can be modelled with a constant, linear, quadratic, or exponential equation (See Supplementary Methods section, SI). Each denaturation curve can have local or shared baseline parameters.

The baseline dependence on denaturant concentration is only considered if the intercepts terms are shared (See Figure 1). In that case, it is assumed to be linear, and a scaling term, for each denaturation curve, can be included in the fitting.

#### ASSUMPTIONS OF THE FITTING MODEL

The fitting model used by CheMelt rests on several assumptions: The protein under study must be a monomer, the unfolding should follow a reversible two-state process. The heat capacity of unfolding Δ*C*_p_ and *m*-value are considered to be independent from temperature and denaturant concentration. Then, the background signal must be negligible (or constant) and that the folded and unfolded states produce a different signal. In this regard, the ensemble of thermally-induced and denaturant-induced unfolded states are considered to be equal. Furthermore, the denaturant contribution to the free energy of unfolding for chemical denaturation is based on the empirical Linear Extrapolation Model (Greene and Pace 1974). These assumptions are not likely to be less precise in the presence of residual structure in the unfolded state but remain the best assumption for most proteins..

## DATA & CODE AVAILABILITY

CheMelt can be used as an online app, Docker image, R shiny application, or Python package (*pychemelt*). The online tool can be accessed at https://spc.embl-hamburg.de/app/chemelt. The Docker image is available at https://hub.docker.com/r/emblspc/chemelt_espc. The R shiny application code is available at https://github.com/SPC-Facility-EMBL-Hamburg/thermochemicalDenaturationApp. The python package *pychemelt* code is available at https://github.com/SPC-Facility-EMBL-Hamburg/pychemelt. Jupyter notebooks with 1) fittings of ACBP and DNAJB6b, 2) fittings of simulated data, and 3) fittings of the hyperstable proteins presented in this manuscript available at https://github.com/SPC-Facility-EMBL-Hamburg/pychemelt_analyses. All experimental data contained in the manuscript is available in FigShare (10.6084/m9.figshare.31889308).

## ACKNOWLEDGEMENTS

This project has received funding from the European Union’s Horizon 2020 research and innovation programme under the Marie Skłodowska-Curie grant agreement No 945405. The work is funded by a grant from the Danish National Research Foundation awarded to the Center for Proteins in Memory (PROMEMO) (DNRF133) and by an instrument supported by Carlsberg Foundation grant (CF21-0164) to M.K. at the Biophysics & Biochemistry Core Facility, Aarhus University, DK. Osvaldo Burastero is funded by the ARISE Fellowship (EMBL and the Marie Skłodowska-Curie Actions). Vili Lampinen and Francisca Pinheiro are funded by Lundbeckfonden (R483-2024-1763 & R449-2023-1396). We thank Kübra Fatima Eroglu for her help in preparing the GluN1 binder proteins, and Jeremias Widmann and Prof. Daniel Otzen for help in doing the circular dichroism measurements.

## USAGE OF LARGE LANGUAGE MODELS

ChatGPT (https://chat.openai.com/) and Github Copilot (https://github.com/features/copilot) were used for coding assistance. The authors take full responsibility for the manuscript content and code.

## SUPPORTING INFORMATION

The supplementary information contains eight figures, three tables and supplementary text describing the theory behind the fitting equations and practical instructions for use.

[Link to Supplementary Information](https://docs.google.com/document/d/1YjgaL8S9vyhEmfZTOgcJsUoPvfTKCWivknP1JwOXeAA/edit?tab=t.0)

