## Supplemental information for "Global analysis of thermal and chemical denaturation using CheMelt: Thermodynamic dissection of highly thermostable *de novo* designed proteins"

### Index of supplementary information:

#### Supplementary text

Theoretical description of the fitting models used by CheMelt.

#### Supplementary figures

**Figure S1:** Comparison of quadratic and exponential unfolded baseline fitting methods for experimental data using ACBP thermal unfolding data.

**Figure S2:** Comparison of quadratic and exponential unfolded baseline fitting methods for experimental data using DNAJB6b thermal unfolding data.

**Figure S3:** Experimental data and global fitting of ACBP unfolding curves.

**Figure S4:** Experimental data and global fitting of DNAJB6b unfolding curves.

**Figure S5:** Simulation and global fitting of hypothetical unfolding curves.

**Figure S6:** Comparison of aromatic residues in proteins with and without a fluorescence-observable unfolding transition.

**Figure S7:** Circular dichroism measurements of binders that did not show signal shift in nanoDSF.

**Figure S8:** Refolding of titered proteins upon cooling.

**Figure S9:** Lack of reversibility suggests that G5A fails to refold binder signal shifts upon cooling

**Figure S10:** Fitting of denaturation data for *de novo* binder proteins.

**Figure S11:** Predicted free energy of unfolding for the thirteen fitted unfolding datasets.

**Figure S12.** Shape and fold of *de novo* binders and native proteins.

**Figure S13.** Intrinsic helicity of natural and de novo designed proteins.

**Figure S14.** Thermodynamic parameters of *de novo* binders and native proteins.

**Figure S15:** SDS-PAGE analysis of purified GluN1 binders

#### Supplementary tables:

**Table S1.** Thermodynamic parameters of ACBP generated by different methods.

**Table S2.** Thermodynamic parameters of DNAJB6b generated by different methods.

**Table S3.** Fit statistics of the DNAJB6b dataset.

**Table S4.** Amino acid sequences of GluN1 binders used in this study.

**Table S5.** AlphaFold2 structure prediction scores of the binders in this study.

**Table S6.** Baseline type of the native and unfolded state selected for each binder.

### Supplementary Methods

#### THEORY

##### COMBINED CHEMICAL AND THERMAL DENATURATION

We start by assuming that the protein is a monomer that (un)folds reversibly via a two-state process. The fraction of protein in the native ($f_n$) and denatured ($f_u$) states depend on the unfolding equilibrium constant $K$.

$$f_n(K) = (1 + K(T, D))^{-1}$$
$$f_u(K) = 1 - f_n$$

where $T$ is the temperature in Kelvin units and $D$ is the denaturant concentration in molar units. The unfolding equilibrium constant $K$ is related to the Gibbs free energy $\Delta G$.

$$K(T, D) = e^{-\frac{\Delta G(T,D)}{RT}}$$

where R is the gas constant. If only chemical denaturation is considered, $\Delta G$ can be calculated using the Linear Extrapolation Model (Greene and Pace 1974).

$$\Delta G(D) = \Delta G_{H_2O} - m \cdot D$$

where $m$ is an empirical parameter that allows modelling $\Delta G$ with a linear dependence on denaturant concentration. $\Delta G_{H20}$ is the free energy of unfolding in the absence of a denaturant agent. On the other hand, if thermal denaturation is considered, the free energy of unfolding can be calculated as indicated below:

$$\Delta G(T) = \Delta H_m(1 - \frac{T}{T_m}) + \Delta C_p(T \quad - T_m - T ln(\frac{T}{T_m}))$$

where $\Delta H_m$ is the enthalpy of unfolding, $T_m$ is the temperature of melting, and $\Delta C_p$ is the heat capacity of unfolding. The combination of both denaturation methods results in the expression:

$$\Delta G(T, D) = \Delta H(1 - \frac{T}{T_m}) + \Delta C_p(T \quad - T_m - T ln(\frac{T}{T_m})) - m \cdot D$$

##### EQUATION OF THE SIGNAL

The total signal is assumed to be a linear combination of the signal produced by the different protein states weighted by their mole fractions.

$$S(\Delta T, D) = f_n(T, D) \cdot S_n(\Delta T, D) + f_u(T, D) \cdot S_u(\Delta T, D)$$

where $S_n(T, D)$ and $S_u(T, D)$ are functions describing the signal dependence on temperature and denaturant concentration for the native and unfolded states, respectively. Here, the background signal is negligible. In regards to differential scanning fluorimetry data, the dependence on temperature has been previously modelled with a quadratic equation (Hamborg et al. 2020):

$$S_x(\Delta T, D = 0) = a + b\Delta T + c\Delta T^2$$

where $a$ is the intercept, $b$ is the linear term, and $c$ is the quadratic term. $\Delta T$ is the difference between the temperature and a reference temperature (298K). If $c$ is zero, then the signal depends linearly on the temperature. If both $b$ and $c$ are zero, then there is no dependence on temperature. Alternatively, the baselines can be modelled with an exponential equation:

$$S_x(\Delta T, D = 0) = \gamma + \delta e^{-\alpha \cdot \Delta T}$$

where $\gamma$ is the intercept, $\delta$ the pre-exponential term, and $\alpha$ the exponential coefficient. Lastly, the dependence on denaturant concentration can be assumed to be linear.

$$S_x(D, \Delta T = 0) = S_x(\Delta T = 0) + d \cdot D$$

where $S_x$ evaluated at $\Delta T$ equal to zero implies that the intercept terms are shared. Here, $d$ is the slope term for the dependence on denaturant concentration.

#### INPUT FILE

To date, Chemelt can parse several kinds of input files including CSV files with column-formatted data, or files exported by instruments of the following manufacturers: Nanotemper (*Prometheus* and *Panta*), Unchained Labs (*AUNTY* and *UNCLE*), Applied Photophysics (*SUPR-DSF*), and Thermofisher Scientific (*QuantStudio™ 3 PCR System*).

#### FITTING

The algorithm for analysing unfolding data in Chemelt is, in essence, the same as the one proposed by Hamborg *et al.* (Hamborg et al. 2020) with two minor differences. The baselines for the native and unfolded states can be modelled with four types of equations: constant, linear, quadratic, and exponential; and secondly, a scale factor is included to account for variances across measurement positions. A more detailed explanation of the fitting procedure is presented below.

To start, all curves must share the same thermodynamic parameters, but there are different possibilities for the baseline terms. A simple option is to fit individual baseline terms for each curve (local intercept and local slopes). In that case, the signal dependence on temperature is modelled separately for each curve, and there is no baseline dependence on denaturant. This method is the most adequate if data from different techniques (or experimental setups, or instruments) are combined.

Then, a more complex option is to share the slope (or exponential) terms (local intercept and global slopes). Therefore, the number of parameters is reduced, but still all curves have an individual intercept term, and the baseline dependence on denaturant is also not considered. This approach is useful when there is a variance in the signal not explained by the temperature or denaturant concentration. For example, if the starting signal for the exact same sample varies.

Finally, to take into account the linear dependence on denaturant concentration (global intercepts and global slopes), and to solve the problem of the experimental, position dependent, signal, the unfolding curves could be rescaled by forcing the signal to be the same at a certain temperature. For example, one could set all curves to have unit value when the temperature is 90ºC. However, this strategy fails if there are curves where the unfolded state is not completely reached at that temperature, which happens for hyperstable proteins. Moreover, it distorts the dependence on denaturant concentration for the unfolded baseline. For this reason, and to allow a global fitting where all baseline terms are shared, we include a scaling factor.

INTERPRETATION

Fitting thermal and chemical denaturation with the proposed model is an ill-posed problem, where a set of unfolding curves can be explained by different parameters values. In that sense, we recommend discarding an individual fitting if the thermodynamic parameters have high errors, if the parameters are physically unreasonable, or if the model predicts certain phenomena that are not observed, such as denaturation at unexpected temperatures (e.g., 5ºC). Moreover, it could happen that the fitting routine gets trapped in local minima if the initial values are far away from the "true" values. To aid the fitting routine, custom bounds can be given in CheMelt for $\Delta H_m$, $T_m$ and $\Delta C_p$.

### Supplementary Figures

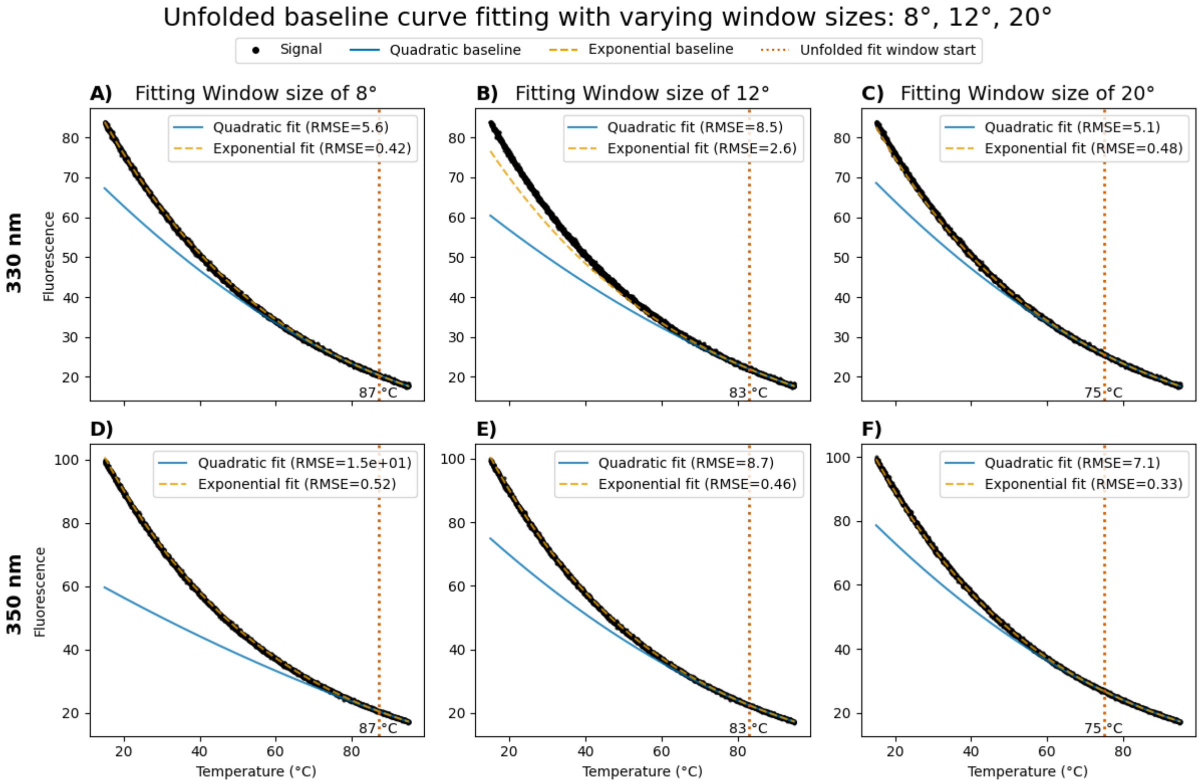


**Figure S1. Comparison of quadratic and exponential unfolded baseline fitting methods for experimental data using ACBP thermal unfolding data**. ACBP thermal unfolding was performed by *Hamborg et al.*[4] between 15 °C and 95 °C of a sample containing 3.01 M guanidium chloride. The data showed no change in curvature and is assumed to correspond to completely unfolded protein. The unfolded baseline was fitted using fluorescence emission signals recorded at 330 nm (A-C) and 350 nm (D-F). The unfolded baseline was fitted as either a quadratic (blue) or exponential (orange) model to the data (black). Fitting to the data was performed with various window sizes where the model was fit to the data higher than the red line. A+D) A fitting window of size 8°C (between 87°C and 95°C) was used. B+E) A fitting window of size 12°C (between 83°C and 95°C) was used. C+F) A fitting window of size 20°C (between 75°C and 95°C) was used. For each fit, the Root Mean Squared Error over the whole dataset is given.

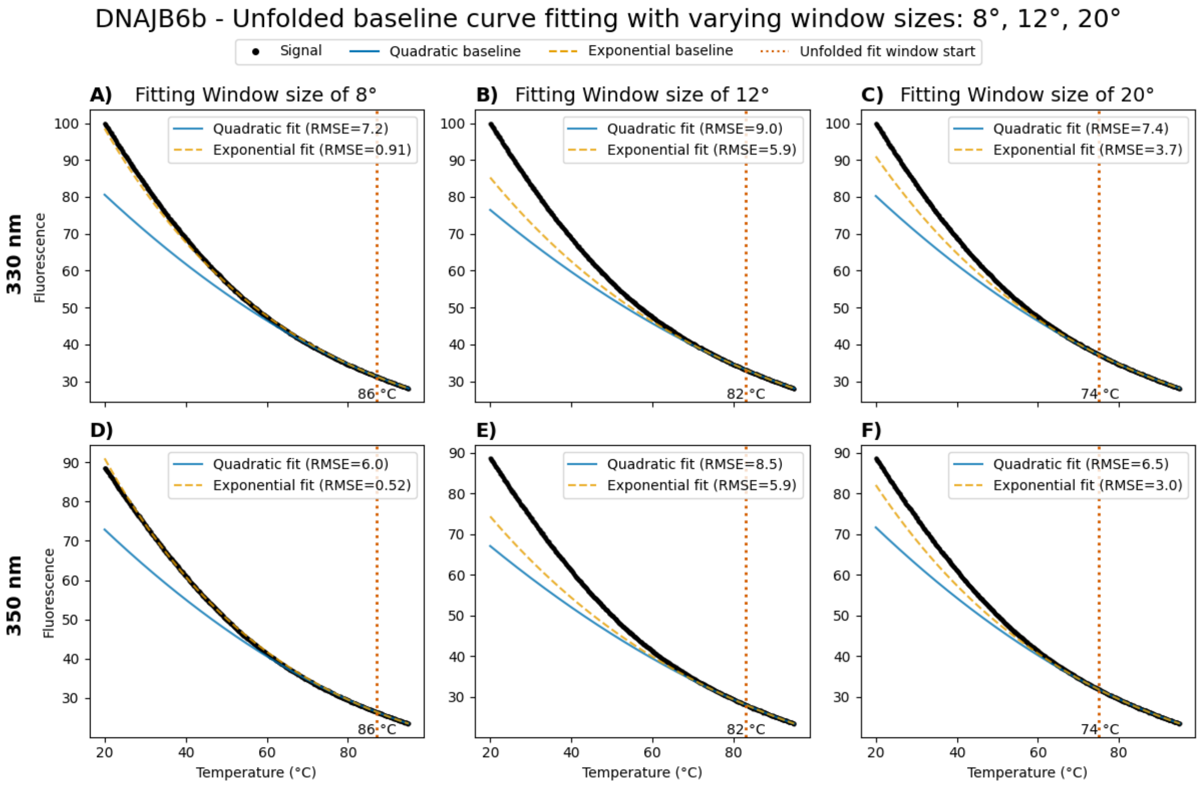


**Figure S2. Comparison of quadratic and exponential unfolded baseline fitting methods for experimental data using DNAJB6b thermal unfolding data**. Thermal unfolding was performed by *Fricke et al.* between 20 °C and 95 °C of a sample containing 4.8 M guanidium chloride. The data showed no change in curvature and is assumed to correspond to completely unfolded protein. The unfolded baseline was fitted using fluorescence emission signals recorded at 330 nm (A-C) and 350 nm (D-F). The unfolded baseline was fitted as either a quadratic (blue) or exponential (orange) model to the data (black). Fitting to the data was performed with various window sizes where the model was fit to the data higher than the red line. A+D) A fitting window of size 8 °C (between 87 °C and 95 °C) was used. B+E). A fitting window of size 12°C (between 83 °C and 95 °C) was used. C+F). A fitting window of size 20 °C (between 75 °C and 95 °C) was used. For each fit, the Root Mean Squared Error over the whole dataset is given.

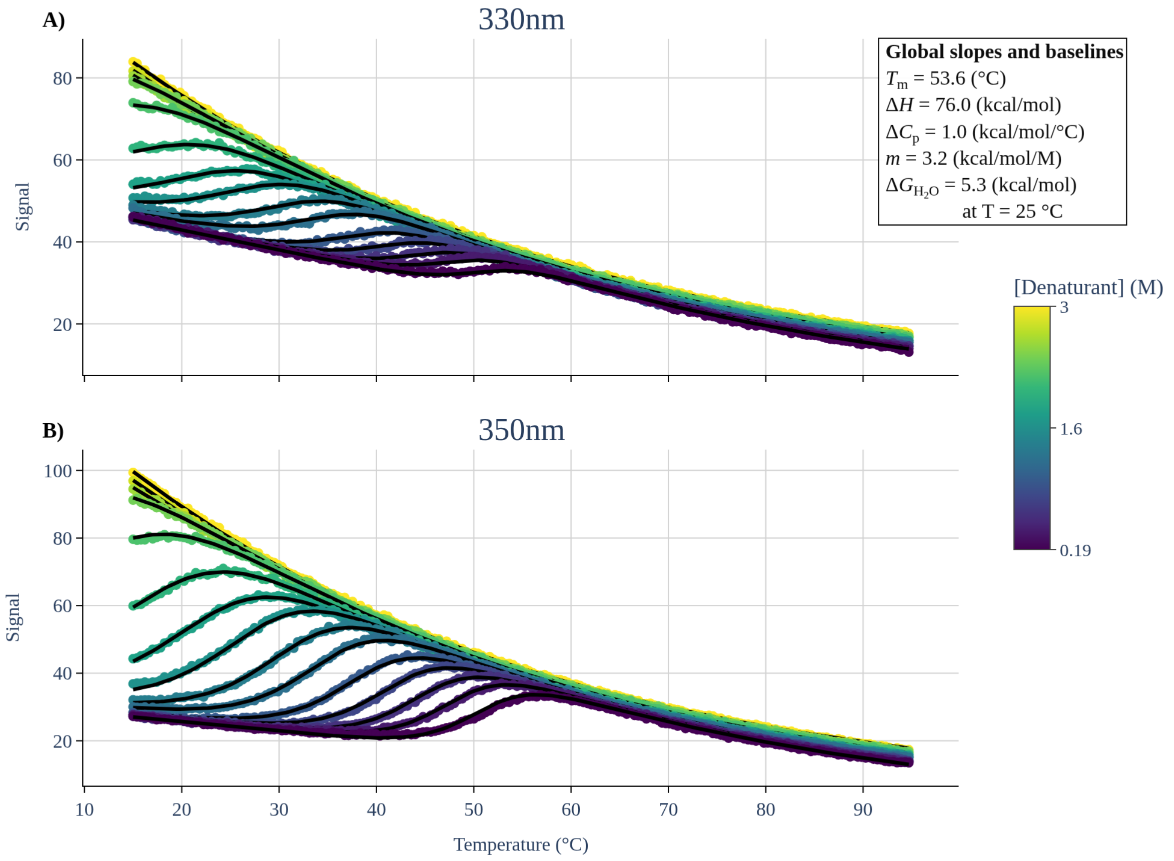


**Figure S3. Experimental data and global fitting of ACBP unfolding curves**. ACBP thermal unfolding was performed by *Hamborg et al.* between 15 °C and 95 °C of samples containing guanidium chloride (GdmCl) at different concentrations between 0.19 M and 3.01 M in approximately 0.2 M steps. Unfolding transitions were fitted using fluorescence emission signals recorded at 330 nm (A) and 350 nm (B). Fitting was performed using a linear function for the native-state baseline and an exponential function for the unfolded-state baseline, with globally shared slope and baseline parameters. A concentration-dependent scaling factor was applied for each GdmCl concentration. Scatter plots show the experimental data, with color coding indicating the GdmCl concentration. Solid black lines represent the global fits to the unfolding model at each denaturant concentration. The resulting thermodynamic parameters melting temperature ($T_m$), enthalpy change ($\Delta H_m$), heat capacity change ($\Delta C_p$), $m$-value, and predicted Gibbs free energy change in water ($\Delta G_{H2O}$) are summarised on the right legend.

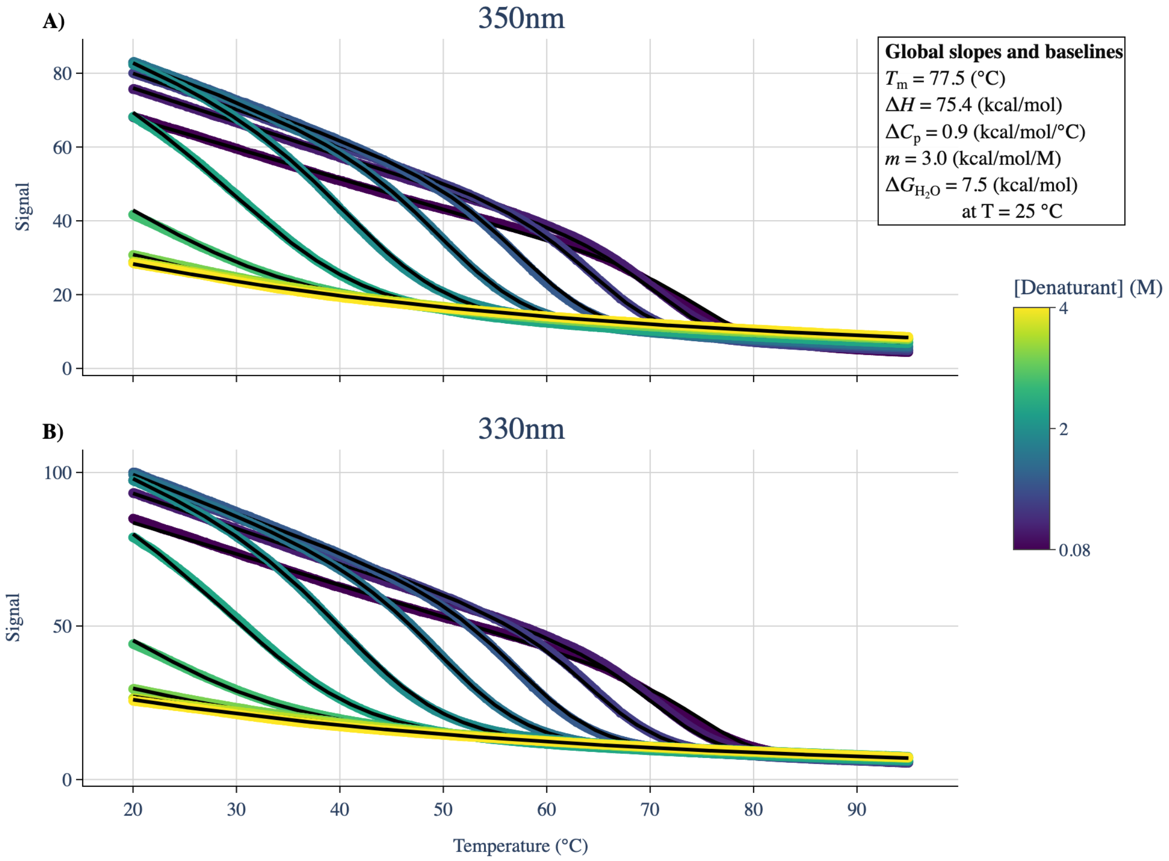


**Figure S4. Experimental data and global fitting of DNAJB6b unfolding curves**. Thermal unfolding was performed by Fricke *et al.* between 20 °C and 95 °C of samples containing guanidium chloride (GdmCl) at different concentrations between 0.08 M and 4 M. Unfolding transitions were fitted using fluorescence emission signals recorded at 330 nm (A) and 350 nm (B). Fitting was performed using a linear function for the native-state baseline and an exponential function for the unfolded-state baseline, with globally shared slope and baseline parameters. A concentration-dependent scaling factor was applied for each GdmCl concentration. Scatter plots show the experimental data, with color coding indicating the GdmCl concentration. Solid black lines represent the global fits to the unfolding model at each denaturant concentration. The resulting thermodynamic parameters melting temperature ($T_m$), enthalpy change ($\Delta H_m$), heat capacity change ($\Delta C_p$), *m*-value, and predicted Gibbs free energy change in water ($\Delta G_{H2O}$) are summarised on the right legend.

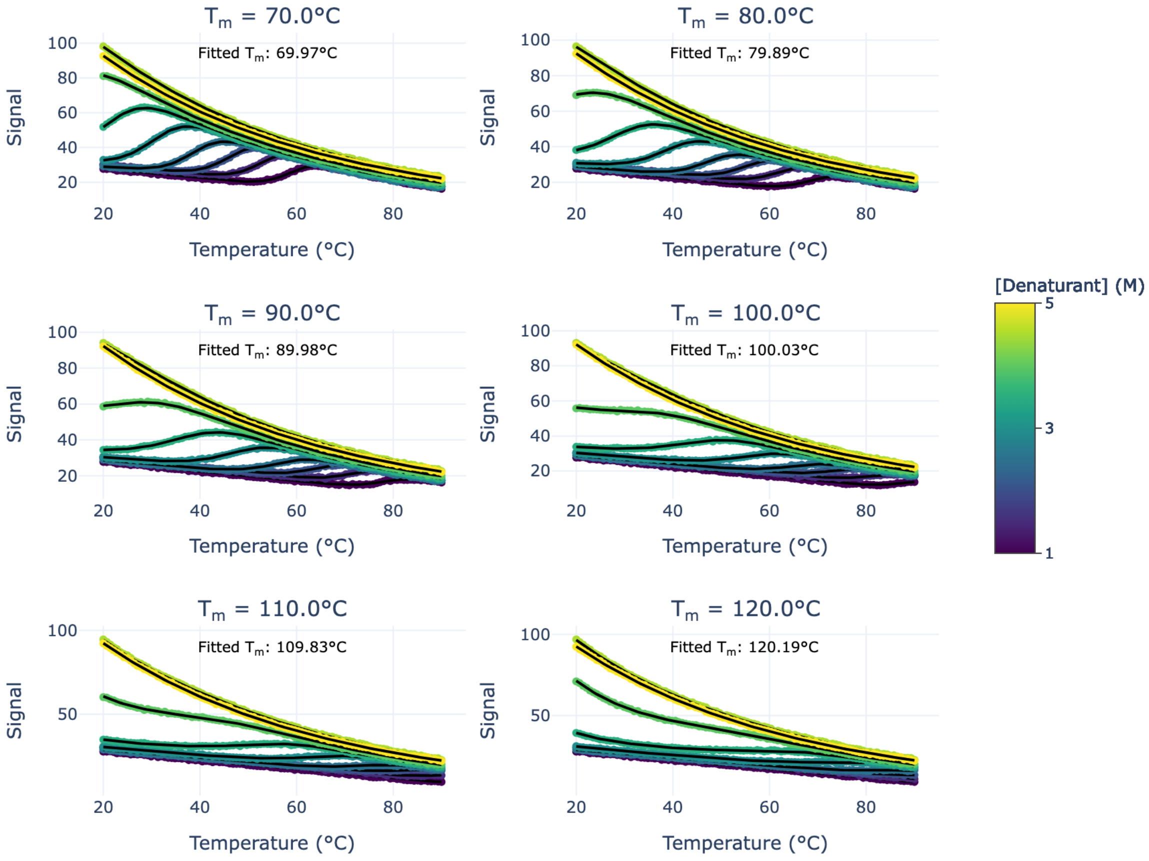


**Figure S5. Simulation and global fitting of hypothetical unfolding curves.** Each panel shares the same parameters for $\Delta H_m$ (100 kcal·mol$^{-1}$), $\Delta C_p$ (1 kcal·mol$^{-1}$·K$^{-1}$), and $m$ (3 kcal·mol$^{-1}$·M$^{-1}$). The folded state baseline follows a linear dependence on temperature ($\Delta T$) and denaturant concentration (D): $S_n(\Delta T, D) = 2.5D + 24 - 0.27\Delta T$. The unfolded state baseline follows an exponential dependence on temperature and linear dependence on denaturant concentration: $S_u(\Delta T, D) = 1.6D - 4 + 80.5e^{-0.0224\Delta T}$. The simulated temperature ranges from 20 to 90°C, while the simulated denaturant concentrations range from 1 to 5 M at 0.5 M steps. Two types of errors were included. Gaussian noise with a standard deviation of 0.25 was added to each data point, and each unfolding curve was scaled by a factor randomly drawn from a uniform distribution between 0.9 and 1.1. The fitted $T_m$ is shown in the plot, while the fitted values for $\Delta H_m$, $\Delta C_p$, and $m$ deviated always less than 1% from the simulated values.

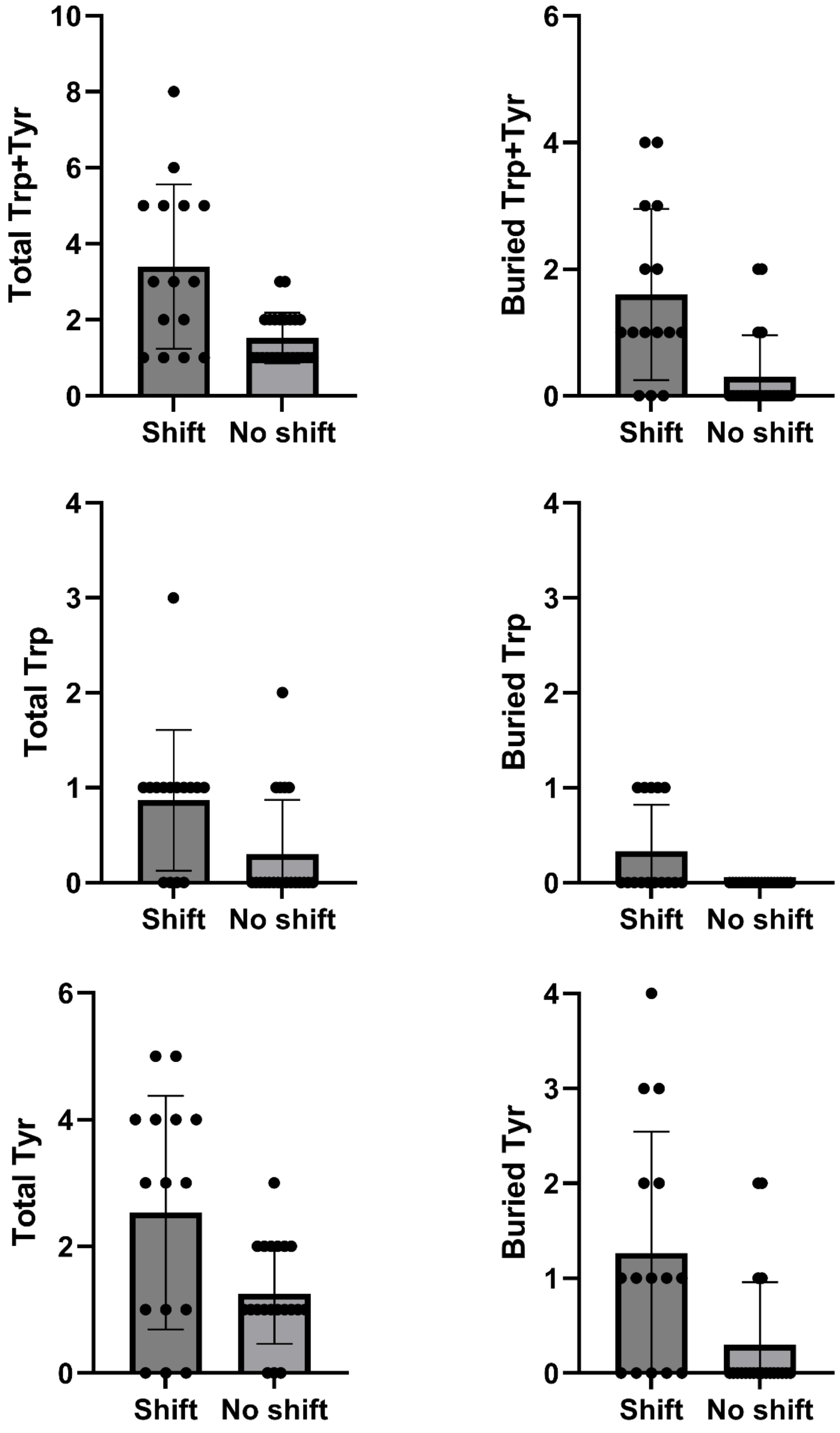


**Figure S6 Comparison of aromatic residues in proteins with and without a fluorescence-observable unfolding transition.** The number of tyrosine and tryptophan residues in each screened *de novo* binder is shown here, with the binders classified to groups with or without a signal shift in nanoDSF measurements. The AlphaFold2 predicted structure of each binder was inspected with PyMol, and each tyrosine and tryptophan was classified to be surface exposed when the exposed area was more than 8.5 Å$^2$.

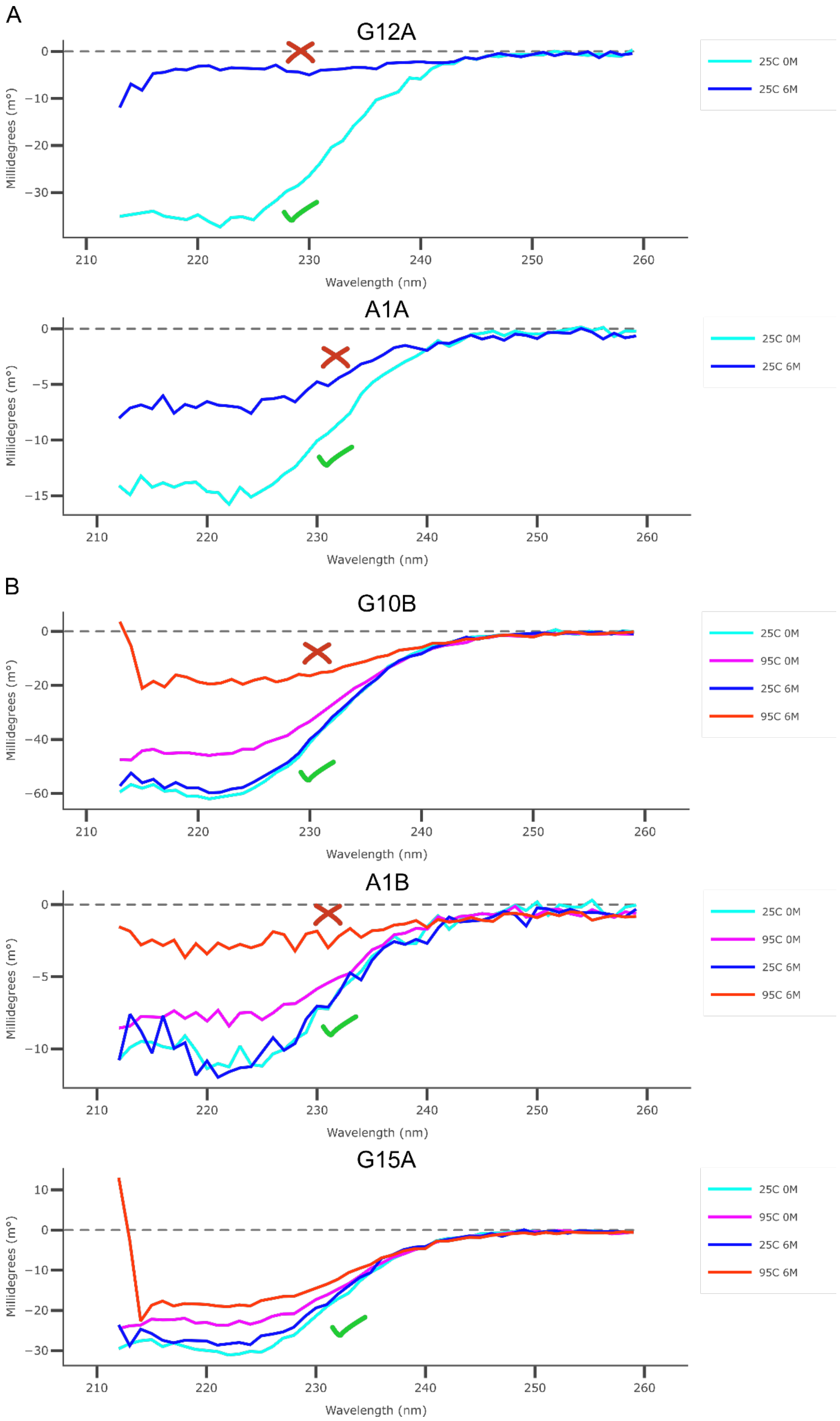


**Figure S7. Circular dichroism measurements of binders that did not show signal shift in nanoDSF.** A) G12A and A1A were measured at 25 °C in PBS and in PBS with 6 M guanidine hydrochloride. B) G10B, A1B, and G15A were measured at 25 °C and 95 °C in the same buffers. All binders but G15A show a condition where the low signal around 220 nm (associated with alpha helix) is mostly lost (indicated with a cross in figure). The graphs were prepared with the ChiraKit web tool.

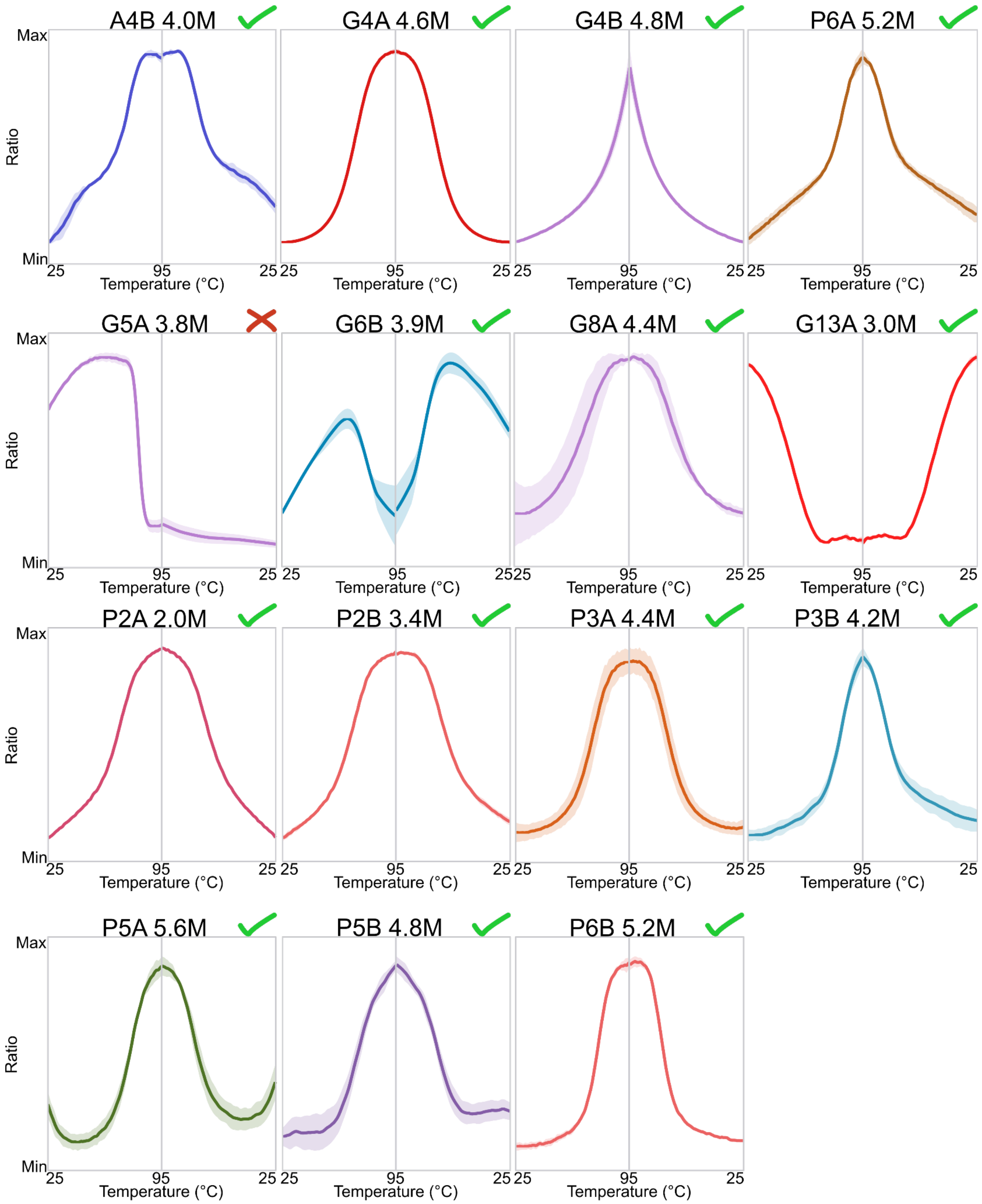


**Figure S8. Refolding of titered proteins upon cooling.** All measurements shown are averages of two independent measurements at a guanidinium chloride concentration with the clearest signal shift. For each protein, the shape of 350 nm /330 nm fluorescence ratio vs temperature is shown. The graph for heating from 25 to 95 °C is shown to the right of the graph for cooling back to 25 °C. The figures were prepared with Prometheus Panta Analysis software.

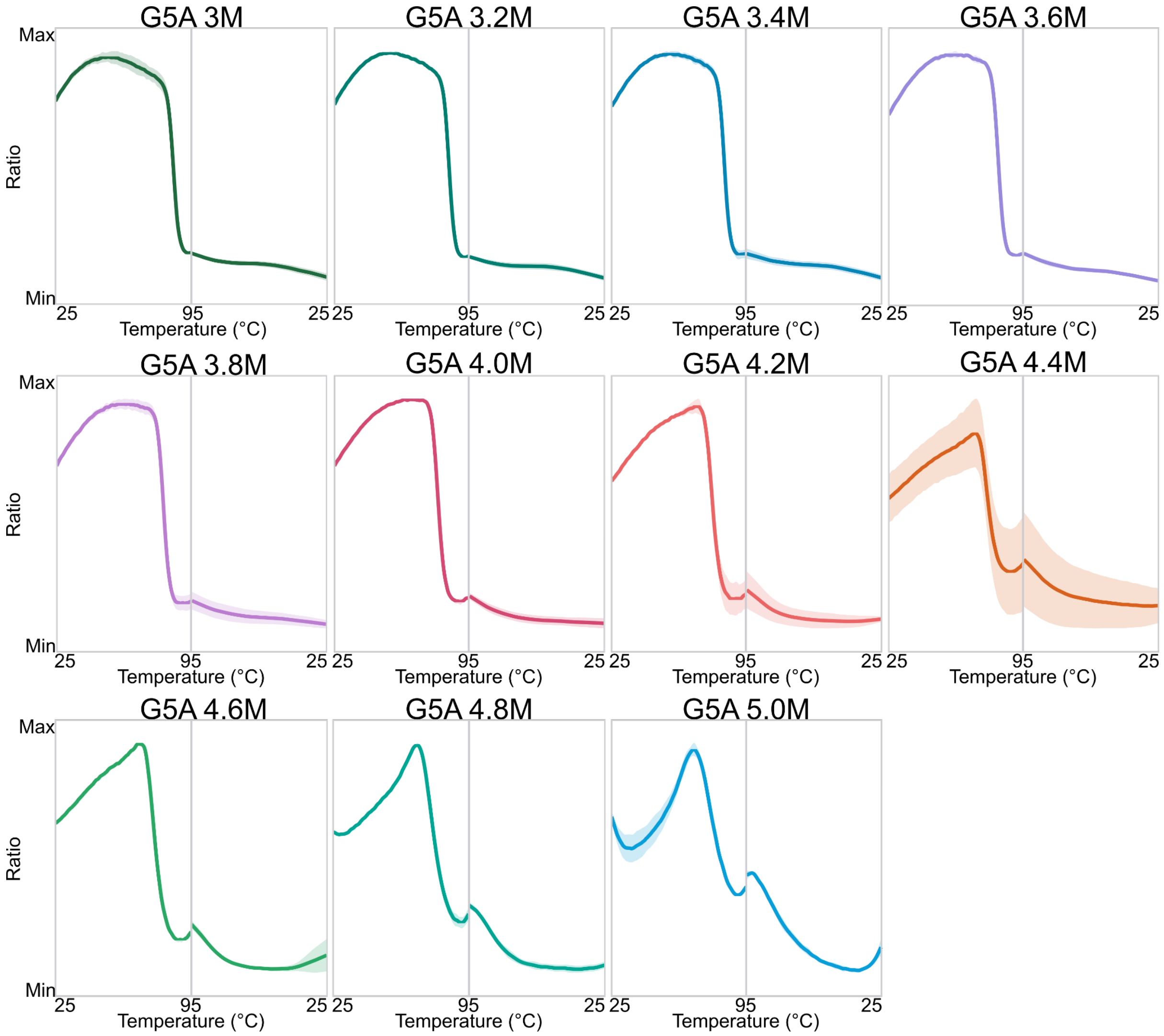


**Figure S9. Lack of reversibility suggests that G5A does not refold upon cooling.** All measurements shown are averages of two independent measurements. For each protein, the shape of 350 nm /330 nm fluorescence ratio vs temperature is shown. The graph for heating from 25 to 95 °C is shown to the right of the graph for cooling back to 25 °C. The figures were prepared with Prometheus Panta analysis software.

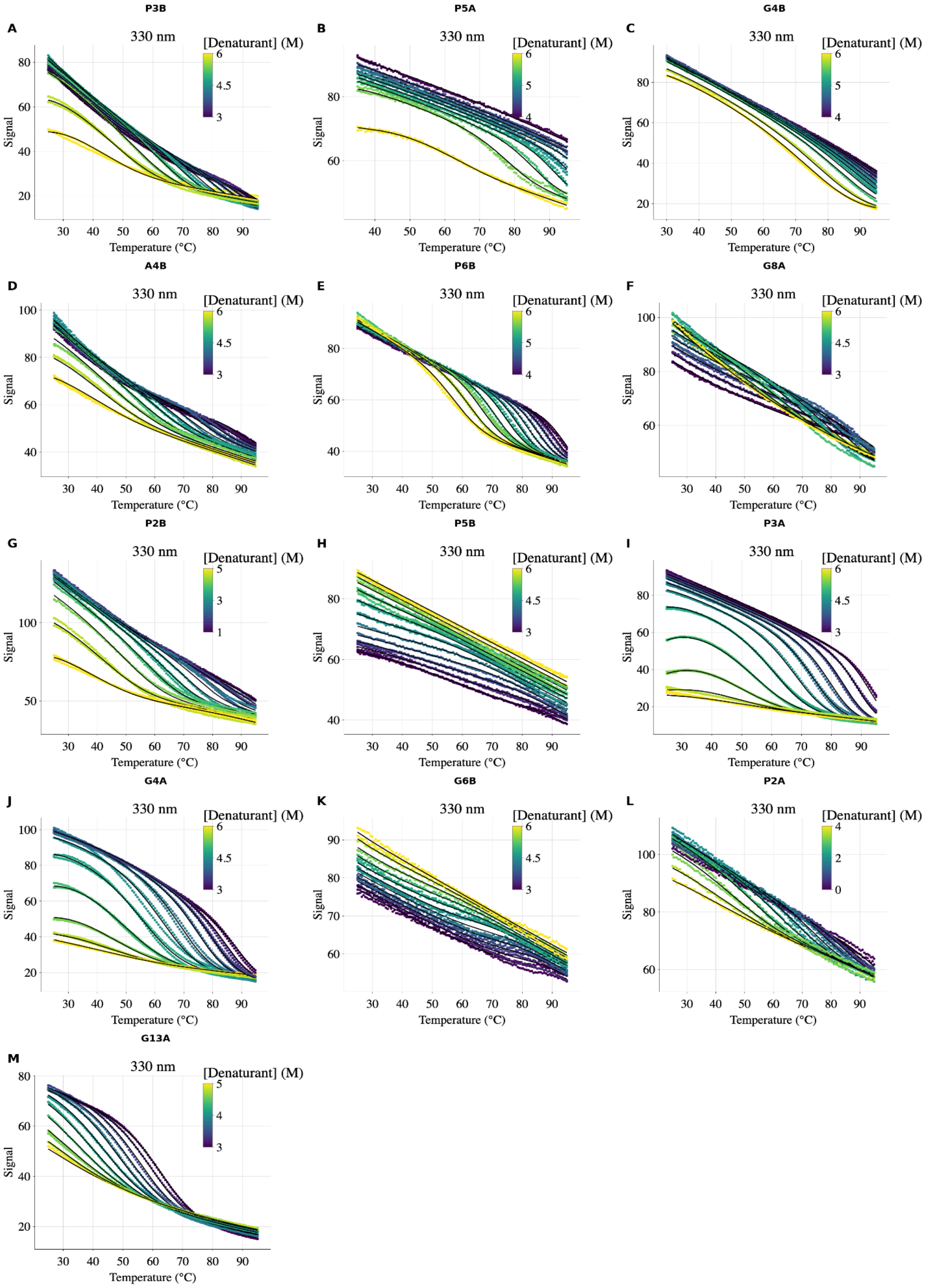


**Figure S10.** Fitting of denaturation data for *de novo* binder proteins. The model was fitted to the 330 nm fluorescence intensity signal for each titered protein. The figure shows the fitted curves after normalisation by the fitted scaled factor to improve clarity. Scatter plots show the experimental data, with color coding (viridis scale) indicating the GdmCl concentration. Solid black lines represent the global fits to the unfolding model at each denaturant concentration.

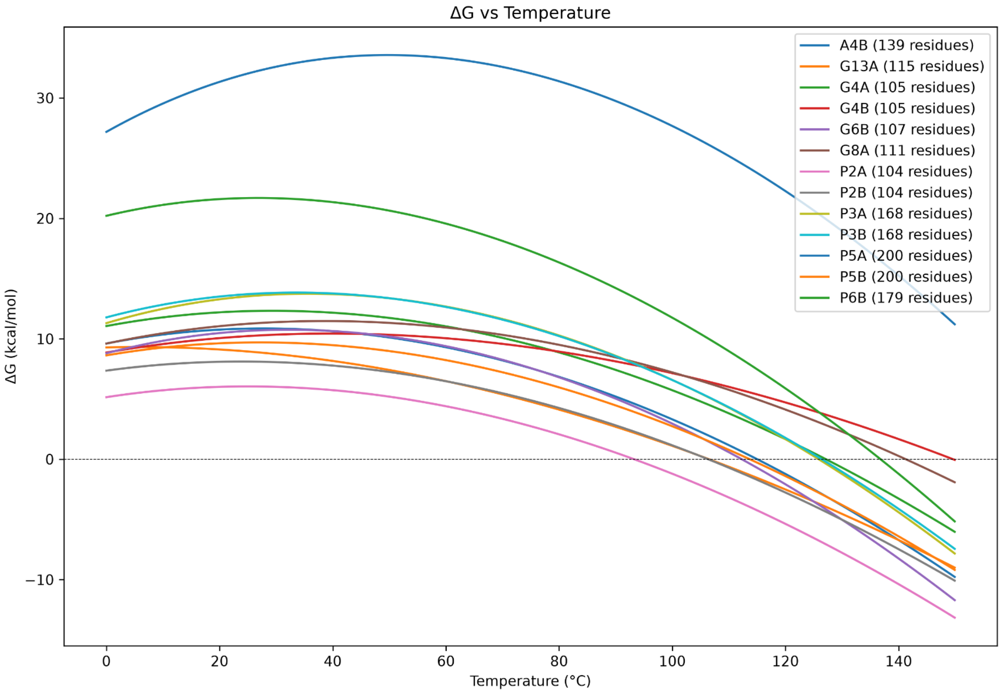


**Figure S11.** Predicted free energy of unfolding as a function of temperature for the thirteen fitted unfolding datasets.

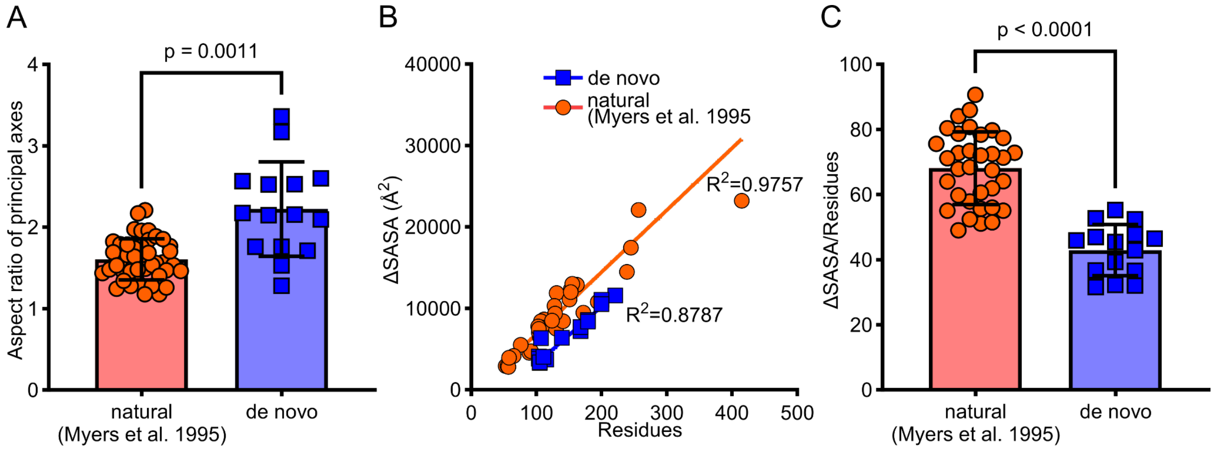


**Figure S12. Shape and fold of *de novo* binders and native proteins.** A) The aspect ratio of principal axes was calculated in regards to the center of mass of each protein model. A value of 1 would represent a perfect sphere, while deviation from this shape increases the value. B–C) Change in surface accessible surface area upon unfolding (ΔSASA) was estimated using the minimum area per residue in the unfolded state (Creamer et al., 1997) and the Shrake–Rupley algorithm for the folded model (Shrake & Rupley, 1973). The native protein models were fetched from RCSB PDB based on the identifiers in Myers et al., 1995. The means of the two groups were compared using Welch's t test (A) and unpaired t test (C).

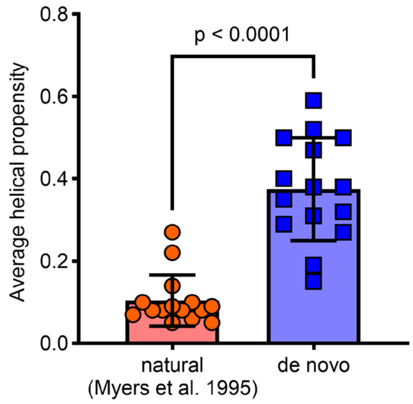


**Figure S13. Intrinsic helicity of natural and de novo designed proteins.** The average helical propensity was estimated for 15 *de novo* proteins showing a measurable unfolding transition and the 15 smallest proteins in the Myers et al. 1995 dataset using ABSEIL. The means of the two groups were compared using Welch's t test.

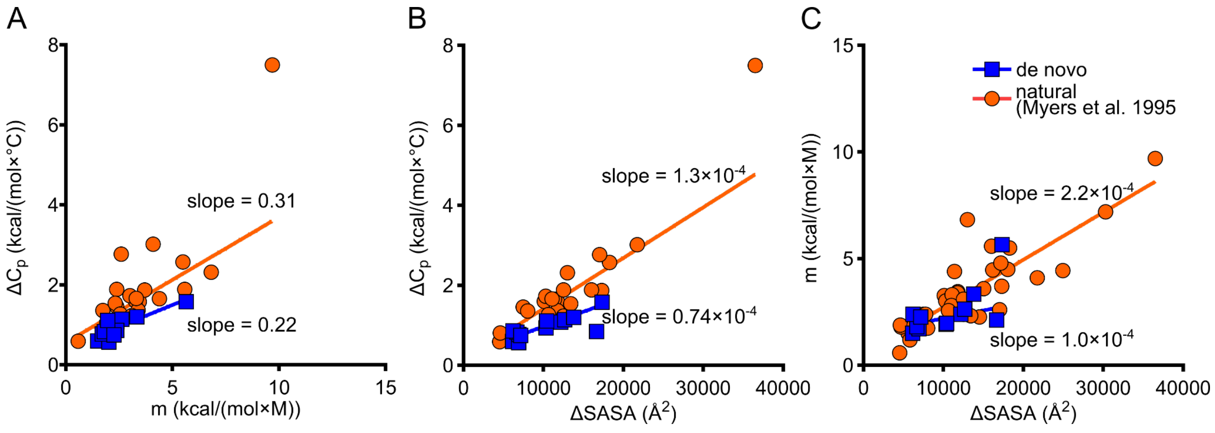


**Figure S14. Thermodynamic parameters of *de novo* binders and native proteins.** The list of native proteins and their thermodynamic parameters were extracted from Myers *et al.* 1995 (Myers et al., 1995). B–C) Change in surface accessible surface area upon unfolding (ΔSASA) was estimated using the maximum area per residue in the unfolded state (Creamer et al., 1997) and the Shrake–Rupley algorithm for the folded model (Shrake & Rupley, 1973). Compare to Figure 6 for an identical analysis using minimum area, which results in smaller ΔSASA. The lines show linear regression of the data.

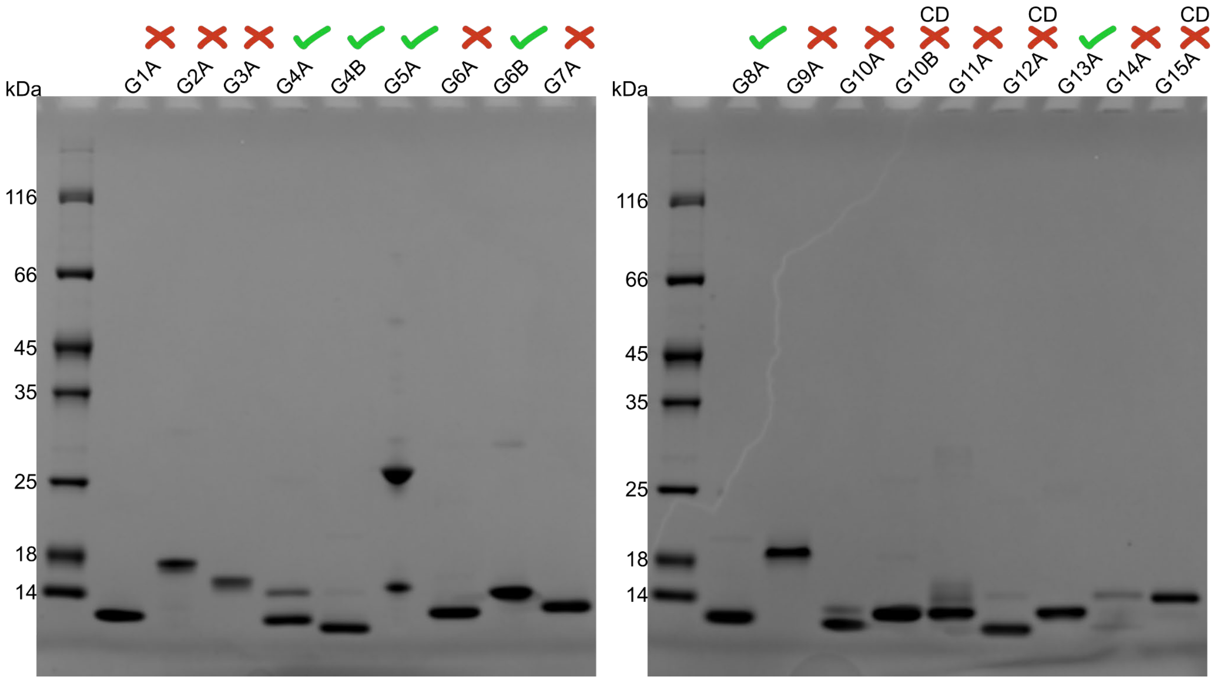


**Figure S15. SDS-PAGE analysis of purified GluN1 binders**. The red crosses indicate the binders that did not show a signal shift in nanoDSF, and CD marks the binders that were analyzed by circular dichroism spectroscopy. Approximately 1 µg of each binder was loaded in each well based on their concentration by absorbance at 280 nm.

### Supplementary Tables

**Table S1. Thermodynamic parameters of ACBP generated by different methods.** CheMelt derived fits are shown in Figure S2. The values from Hamborg *et al.* are the averages and standard deviations of three independent experiments. No replicates were performed for CheMelt.

| **Reference** | $T_m$ (°C) | $\Delta H_m$ (kcal/mol) | $\Delta C_p$ (kcal/mol/°C) | ***m*-value (kcal/mol/M)** | $\Delta G_{H2O}$ (kcal/mol) |
|---|---|---|---|---|---|
| This work* | $53.61^{+0.07}_{-0.07}$ | $75.96^{+0.58}_{-0.58}$ | $1.03^{+0.02}_{-0.02}$ | $3.2^{+0.02}_{-0.02}$ | 5.32 |
| (Hamborg et al. 2020) | 53.05 ± 0.1 | 82.46 ± 3.6 | 1.1 ± 0.14 | 3.54 ± 0.07 | 5.76 ± 0.14 |
| (Teilum et al. 2002)** | - | - | - | 3.51 | 5.98 |

*Subscripts and superscripts provide the 95% confidence interval estimated using the profile likelihood method.

**Teilum *et al.* reported errors, but it is unknown if it corresponds to fitting errors, or standard deviations of independent experiments.

**Table S2. Thermodynamic parameters of DNAJB6b generated by different methods.** CheMelt derived fits are shown in Figure S4.

| **Reference** | $T_m$ **(°C)** | $\Delta H_m$ **(kcal/mol)** | $\Delta C_p$ **(kcal/mol/°C)** | ***m*-value (kcal/mol/M)** |
| --- | --- | --- | --- | --- |
| This work* | $77.53^{+0.01}_{-0.01}$ | $75.42^{+0.01}_{-0.01}$ | $0.93^{+0.00}_{-0.00}$ | $3.00^{+0.00}_{-0.00}$ |
| Fricke *et al.*, 2025 | 77.41 ± 0.04 | 63.2 ± 0.2 | 0.72 ± 0.02 | 2.6 ± 0.07 |

*Subscripts and superscripts provide the 95% confidence interval estimated using the profile likelihood method.

**Table S3. Fit statistics of the DNAJB6b dataset.** The code from Fricke *et al.*, is available at **https://data.dtu.dk/articles/code/Fitting_of_thermal_and_chemical_denaturation/27679716**

| | Fricke *et al.*, 2025 | This work |
|---|---|---|
| data points | 20328 | 20328 |
| variables* | 18 | 28 |
| chi-square | 92545172.5 | 3274 |
| reduced chi-square | 4557 | 0.16 |
| Akaike information criteria | 171268 | -37062 |
| Bayesian information criteria | 171411 | -36840 |

* The number of variables differs because of the inclusion of a scale factor in CheMelt.

**Table S4.** Amino acid sequences of GluN1 binders in this study.

Color coding: 6xHis-tag, Thrombin cleavage site, Binder

| |
|---|
| **GluN1 binder 1A** |
| MGSSHHHHHHSSGLVPRGSHMYEFLERARRVLETLARLEELLARVDPEAAERAAELREEVEEVIERVEEEGDVEEAEERLEELRERALELIREARERL |
| **GluN1 binder 2A** |
| MGSSHHHHHHSSGLVPRGSHMMEAEIAALRAAIEAAADDPVRRAVLLAQLALLLLQAGREEEADAAFLEFLDLMASLSPEERAAAFRAILDALLAWPAADIWRFFLRFAEVAVRAGRVAELLDLFRALLEAARARAAETGTPAAATLLRIVEAILRIVEEHLARLEAA |
| **GluN1 binder 3A** |
| MGSSHHHHHHSSGLVPRGSHMMKVHVADDVEEALAILLETKDEDVLVLLTRELSPEEAIAAFEEAARLMKEKAAGTRKVLVLLVPDAAASREPLAALKERLETEMGFAAEIRELPSGRVVAVLAYAPPEEAEEAKELAREVAERVAALVERILAER |
| **GluN1 binder 4A** |
| MGSSHHHHHHSSGLVPRGSHMELVELTPEQEAALEADLAARLKAAGLDAATDEFVAIWRRGLAAGDIAGTVAELLARAKEIGEEYVPTVREILLAIVRRLLELAA |
| **GluN1 binder 4B** |
| MGSSHHHHHHSSGLVPRGSHMEEVELTPEQLEALLEALRAALEAAGLTAHVEEFVAIWREGLAAGDIAGTVERLLALAEEIGPEHVPAVRAILLAVVRRLLELAA |
| **GluN1 binder 5A** |
| MGSSHHHHHHSSGLVPRGSHMAEEEAARREALLARARAGEEIPPEDWRDVMLAGLAADPATRSVYDLIEKYAELVGRYRAALSALVERAVERFFEGEEVEIEALVLPADILANPAVTDAELRAFLEALTARLEAALALDPAEHEAALRAVLADVFAYAAAQGVRVVVAVWSRRVAATLYVADLVVDVAKANPDVTPEMLTIRVFNREAARIAAWVARRLAA |
| **GluN1 binder 6A** |
| MGSSHHHHHHSSGLVPRGSHMELMAQARAIAEAAGSVFMLAALDEAERARSLPMEELRKRLAEAAEALAADPSHLAAYIRASTVYILHSSPAEAREAHARLLAALAAARARAGPERAALLDRLAAVHEAILELKEAADAA |
| **GluN1 binder 6B** |
| MGSSHHHHHHSSGLVPRGSHMYSEAERQLEIIRRVLLRMLEAGERERLAEIVNAIREVLGLPKVSLEEVTEELIEELVEEVRRRPEALPAVRAIVLEALRRMLAEAE |
| **GluN1 binder 7A** |
| MGSSHHHHHHSSGLVPRGSHMYSPRLQALLAETQRLLAAARAGTLETREREARRLRELAREAEAAAAEVAADPALRAELEAFARRVRALARALEELVAAERA |
| **GluN1 binder 8A** |
| MGSSHHHHHHSSGLVPRGSHMMTPEELAAARAAFLARAREAAALDPAERMAALRELLEEARAWFLAVAADPENRLAAVEAFLAFLREVVALAPEARPLALEVLREVRAALA |

**GluN1 binder 9A**

MGSSHHHHHHSSGLVPRGSHMELMAQARAIAEAAGSVFMLAALDEAERARSLPM
EELRKRLAEAAEALAADPSHLAAYIRASTVYILHSSPAEAREAHARLLAALAAARA
RAGPERAALLDRLAAVHEAILELKEAADAA

**GluN1 binder 10A**

MGSSHHHHHHSSGLVPRGSHMDLMERARAIAEEAGSVFMLAALLEAEKDKSLPM
EEIKKRYEEAKAALKADPSKLPEFIRATTVYILHSSLEEAEAATKELLAALDAAAA
QASPERRALLDRVREVVLRILELKKEAAEK

**GluN1 binder 10B**

MGSSHHHHHHSSGLVPRGSHMMEEEIERAEREILLAVLRELAALLDAGERLTLPEA
RERARALIEERVAELPEEIRERVRERVLARLDALLARWAAE

**GluN1 binder 11A**

MGSSHHHHHHSSGLVPRGSHMYEEREQQLRIALALVERILEALEKGELVEPEVLEH
LLERLKKLGLEEEAKLMEEALELRLKALEDPSTFEENFKKMLEKVKEALELLKKK
A

**GluN1 binder 12A**

MGSSHHHHHHSSGLVPRGSHMEELVELPPEEAARLRAVMEFVRLMDKPLEEMEA
WVAEREETRALMEELLALYKADPAAARARLLAELRAALARAA

**GluN1 binder 13A**

MGSSHHHHHHSSGLVPRGSHMSRAARAARILALIAELEAALADPSLSPAERLAARV
EALTEILELEGEDREEARALAERLAALHAAAIEAGREAEAWAILRAGLAELRAAAL
ELA

**GluN1 binder 14A**

MGSSHHHHHHSSGLVPRGSHMGREEKLKRKREVLIRVRERIRRAYLAEGKEERAE
EVEKELEEFEKEFEKLSLEEQEKLLEEESKKVLEEARRLLEEQE

**GluN1 binder 15A**

MGSSHHHHHHSSGLVPRGSHMMEAEIERLVREILLEVLREILELLDRGEELTPEEAR
ERARELIEELVEKLPEEIREEVLERALEELERLLARWEAE

**Table S5. AlphaFold2 structure prediction scores of the binders in this study.**

| Binder | Average predicted alignment error (Å) |
|---|---|
| **A1A** | 3.436 |
| **A1B** | 3.441 |
| **A2A** | 3.671 |
| **A3B** | 3.811 |
| **A4A** | 3.82 |
| **A4B** | 3.925 |
| **G1A** | 2.379 |
| **G2A** | 2.703 |
| **G3A** | 3.628 |
| **G4A** | 2.575 |
| **G4B** | 2.76 |
| **G5A** | 3.021 |
| **G6A** | 2.302 |
| **G6B** | 2.733 |
| **G7A** | 2.657 |
| **G8A** | 2.528 |
| **G9A** | 2.573 |
| **G10A** | 2.884 |
| **G10B** | 2.985 |
| **G11A** | 3.284 |
| **G12A** | 3.516 |
| **G13A** | 2.523 |
| **G14A** | 2.527 |
| **G15A** | 3.019 |
| **P1A** | 3.881 |
| **P1B** | 3.975 |
| **P2A** | 3.884 |
| **P2B** | 4.004 |
| **P3A** | 3.865 |
| **P3B** | 3.903 |
| **P4B** | 3.93 |
| **P5A** | 4.671 |
| **P5B** | 4.807 |
| **P6A** | 6.114 |
| **P6B** | 2.827 |

**Table S6. Baseline type of the native and unfolded state selected for each binder.**

| Name | Native Baseline | Unfolded Baseline |
|---|---|---|
| **P5A** | quadratic | linear |
| **G4B** | linear | linear |
| **G8A** | exponential | exponential |
| **P6B** | quadratic | linear |
| **G4A** | quadratic | quadratic |
| **P3B** | quadratic | quadratic |
| **P3A** | quadratic | linear |
| **A4B** | quadratic | linear |
| **P5B** | quadratic | linear |
| **G6B** | exponential | quadratic |
| **G13A** | quadratic | quadratic |
| **P2B** | exponential | quadratic |

Hamborg, Louise, Emma Wenzel Horsted, Kristoffer Enøe Johansson, Martin Willemoës, Kresten Lindorff-Larsen, and Kaare Teilum. 2020. "Global Analysis of Protein Stability by Temperature and Chemical Denaturation." *Analytical Biochemistry* 605 (September): 113863.

Teilum, Kaare, Birthe B. Kragelund, and Flemming M. Poulsen. 2002. "Transient Structure Formation in Unfolded Acyl-Coenzyme A-Binding Protein Observed by Site-Directed Spin Labelling." *Journal of Molecular Biology* 324 (2): 349–357.
